# Clinical performance evaluation of a tiling amplicon panel for whole genome sequencing of respiratory syncytial virus

**DOI:** 10.1101/2025.01.26.634892

**Authors:** B. Ethan Nunley, Amelia Weixler, Hyeong Geon Kim, Hong Xie, Jaydee Sereewit, Pooneh Hajian, Sean Ellis, Margaret G. Mills, Ailyn C. Pérez-Osorio, Stephanie Goya, Jolene Gov, Rebecca Dewar, Goncalo Fernandes, Kate E. Templeton, Daniel M. Maloney, Alexander L. Greninger, Pavitra Roychoudhury

## Abstract

Accurate genomic characterization of respiratory syncytial virus (RSV) is crucial for studies of epidemiology and viral evolution, and monitoring potential escape from newly authorized vaccines and antivirals. We adapted a viral whole genome tiling amplicon panel (UW-ARTIC) and developed a custom bioinformatic pipeline for high-throughput, cost-effective sequencing of RSV-A and RSV-B. We established genome acceptability criteria and determined the performance characteristics of the panel including assay sensitivity, specificity, breadth of genome recovery, accuracy, and precision using contrived and remnant clinical specimens. High-quality genomes (>95% genome completeness; >500X and >1000X average depth for whole genome and fusion gene respectively) were recovered from samples with Ct ≤ 30 (∼594 and 2,004 copies per reaction for RSV-A and RSV-B respectively). Minor variants were accurately identified in sample mixtures of 5:95 and higher. The assay showed high accuracy when compared against Sanger, shotgun metagenomic, and hybridization capture-based sequencing; and high repeatability and reproducibility. The UW-ARTIC RSV panel has utility in genomic surveillance, clinical and research applications. It has been used to generate FDA-reportable data for clinical trials of RSV antiviral products, with robust performance characteristics in samples from around the globe from as recently as the 2023/24 season. Continued genomic surveillance and future updates to primer sets will be essential for continued recovery of genomes as RSV continues to evolve.

## Introduction

Human respiratory syncytial virus (RSV) is a leading cause of severe respiratory illness in infants, older adults, and immunocompromised individuals [1] resulting in over 30 million cases of acute lower respiratory infection annually around the world [2,3]. The approval of new RSV vaccines for pregnant persons and older adults, and monoclonal antibodies like nirsevimab for passive immunization of infants, mark important steps towards reducing RSV-associated disease burden [4]. Additionally, multiple antiviral agents in various stages of development provide promising therapeutic options for managing RSV infections. Tracking the evolution and spread of RSV using genomic sequencing is crucial for identifying variants with potential to cause more severe disease or prophylactic therapeutic resistance [5].

Tiling amplicon panels are a cost-effective and efficient approach for viral whole genome sequencing for both genomic surveillance and clinical applications. During the COVID-19 pandemic, multiple tiling amplicon workflows became available for SARS-CoV-2 whole genome sequencing including the ARTIC nCoV-2019 panel incorporated into the COVIDseq assay (Illumina), NEBNext (New England Biolabs), xGen/Swift Normalase Amplicon Panel (IDT), and the QIAseq Direct kit (QIAGEN), which have since been extended to other pathogens. Many of these workflows can be adapted with plug-and-play primers for pathogens of interest with options for automation using high-throughput liquid handlers that allow surveillance programs to be scaled up rapidly [6].

Here we adapted RSV ARTIC primer sets described previously [7,8] for the recovery of whole RSV-A and RSV-B genomes. These primers were used to validate a sequencing workflow for RSV that leverages existing methods originally designed for SARS-CoV-2 high-throughput sequencing in a clinical laboratory [9]. The amplicon panel used in this validation (UW-ARTIC) consists of 50 pairs of forward and reverse primers that produce amplicons 400-500 bp in length, with separate primer sets for RSV-A and RSV-B.

Our validation methodology was based on available FDA guidance for validation of next-generation sequencing protocols for germline diseases [10], and our prior clinical validation methods for SARS-CoV-2 [9,11]. Contrived and remnant clinical samples were used to determine the assay’s analytical sensitivity, specificity, detection of minor alleles, precision, and accuracy, and genome acceptability criteria were defined to ensure quality of genomes recovered.

## Materials and Methods

### Specimens and RNA extraction

This study used remnant clinical specimens (Copan swabs in UTM) tested at the University of Washington Virology Lab. Use of remnant specimens for this study was approved by the University of Washington Institutional Review Board with a consent waiver (protocol STUDY00010205). Remnant specimens from October 2022 to February 2023 were stored in viral transport media at −80°C until use in this validation. Nucleic acid was extracted from 200µL of each specimen, and eluted to 50µL using the MagNA Pure 96 DNA and Viral NA Small Volume Kit on the MagNA Pure 96 instrument (Roche Diagnostics) according to manufacturer’s instructions, 40 µL of 10^5^ copies/µL EXO RNA was spiked into 5 mL lysis buffer to achieve 100 copy/µL of eluted extraction material concentration in sample as internal control.

### Amplicon panel for RSV whole genome sequencing

The amplicon panel developed for this study was based on the ARTIC RSV primer scheme described previously [7,8]. The original ARTIC RSV-A and RSV-B primer schemes were designed using 6 recent RSV-A and 6 RSV-B sequences and consisted of two sets of 50 primer pairs designed to generate fragments ∼400bp in length. After initial testing, modifications were made to the original design (to primers RSVB_17_RIGHT, RSVB_17_LEFT, and RSVB_45_RIGHT) to accommodate circulating diversity based on Washington State sequences collected up to January 2023 and to prevent amplicon drop-out. The modified primer scheme (UW-ARTIC) consists of two sets of 50 primer pairs for RSV-A and RSV-B respectively (Table S1).

### RT-PCR for typing and quantification

Typing of RSV samples was performed using real time RT-qPCR targeting the polymerase gene with 10 μL of template added to 15 μL of typing PCR master mix prepared using AgPath-ID One-Step RT-PCR reagents (ThermoFisher Scientific, catalog #4387424). Typing PCR master mix consisted of 12.5 μL of AgPath-ID One-Step 2X RT-PCR buffer, 1 μL of AgPath-ID 25x enzyme mix, 0.2 µL RSV typing primer mix containing forward and reverse primers (Table S2) at 50 µM each, 0.06µL RSV probe (Table S2) mix containing RSV-A and RSV-B probes at 50 µM each, 1 μL of EXO3 primer mix containing 3.5 µM forward µM and 7 µM reverse primers (Table S2), 0.22 μL of 10 µM EXO3 Cy5 probe (Table S2), and 0.02 μL of molecular-grade, nuclease-free water. RT-qPCR was performed on ABI 7500 instruments (Thermo Fisher) with the following thermocycling conditions: 50°C for 15 minutes, 95°C for 2 minutes, 40 cycles of 95°C for 15 seconds and 60°C for 1 minute. PCR results were analyzed using Applied Biosystems 7500 Software v2.3 with a baseline start of 3, a baseline end of 15, and a threshold of 40,000 for each dye.

Quantification of RSV RNA in extracted samples was performed using real time RT-qPCR targeting the matrix gene. 10 μL of template was added to 15 μL of qPCR master mix. Quantification master mix consisted of 12.5 μL AgPath-ID One-Step 2x RT-PCR Buffer, 1 μL of AgPath-ID One-Step Enzyme Mix, 0.27 μL RSV primer mix containing forward and reverse primers at 25 µM each, 0.03 μL of RSV FAM probe at 100 µM, 1 μL of EXO2 primer mix (Table S2) containing forward primer at 3.5 µM and reverse primer at 7 µM, and 0.2 μL of 10 µM EXO2 VIC probe (Table S2) per reaction. RT-qPCR was performed on ABI 7500 instruments with the following thermocycling conditions: 50°C for 15 minutes, 95°C for 2 minutes, 40 cycles of 95°C for 15 seconds and 60°C for 1 minute. PCR results were analyzed using Applied Biosystems 7500 Software v2.3 with a baseline start of 3, a baseline end of 15, and a threshold of 0.05 for RSV FAM probe and automatic baseline values and a 0.05 threshold for EXO2 VIC probe.

### Library Preparation, sequencing, and quality control

Sequencing libraries were prepared using the Illumina COVIDseq kit (catalog #20044461) as follows. After extraction, single-stranded cDNA (sscDNA) was created by first priming 8.5 µL of RNA by adding 8.5 µL EPH3 and incubating on a thermal cycler at 65°C for 3 minutes, then adding 8 µL of master mix made with 9 µL First Strand Mix to 1 µL Reverse Transcriptase (extra volume was included in making the master mix to account for pipetting coverage) and incubating under the following conditions: 25°C for 5 minutes, 50°C for 10 minutes, and 80°C for 5 minutes, with a 4°C hold. Amplicon PCR was performed on sscDNA using 20 µL of COVIDseq PCR master mix, made with 15 µL Illumina PCR Mix, 1.7 µL Nuclease-free water, and 4.3 µL of the respective amplicon primer pool mix at 14 µM, but replacing COVIDseq pool 1 and pool 2 amplicon primers with UW-ARTIC primer pool 1 and 2 for either RSV A or RSV B, depending on the samples in the batch. Thermocycler conditions were as follows: 98°C for 3 minutes, 34 cycles of 98°C for 15 seconds and 63°C for 5 minutes, with a 4°C hold. The lid temperature was set to 105°C. Following amplicon PCR, each sample was run on D1000 Tapestation (Agilent, catalog #5067-5582) to determine the presence of amplification, based on the presence of a peak with a normalized sample intensity greater than 100 within Agilent Tapestation Analysis Software v5.1. After QC, 10 µL of PCR product from pool 1 and pool 2 were combined and libraries were prepared using COVIDseq reagents and manufacturer instructions. 30 µL of tagmentation mix, made with 12 µL Tagment Buffer 1, 4 µL Enrichment Bead-Linked Transposomes, and 20 µL nuclease-free water was added to the combined 20 µL of pool 1 and 2 PCR products and incubated for 5 minutes at 55°C. Tagmentation was halted after a 2-minute room temperature incubation with the addition of 10 µL Stop Tagmentation Buffer 2. Tagmented samples were washed twice with 100 µL Tagmentation Wash Buffer and eluted into 40 µL of library PCR master mix consisting of equal parts nuclease-free water and Enrichment PCR Mix. 10 µL of Illumina dual-index adapters were added to each sample and amplified under the following conditions: 72°C for 3 minutes, 98°C for 3 minutes, twelve cycles of 98°C for 20 seconds, 60°C for 30 seconds, and 72°C for 1 minute, with a final extension of 72°C for 3 minutes and a 10°C hold. Following PCR, the 1 µL of up to 12 randomly chosen libraries were run on a 1.2% Agarose 12+1 lane Flashgel (Lonza, catalog #57023) to ensure a visible smear in the 270-750bp range. Libraries were then pooled in equal volumes and a 0.8x cleanup was performed using Illumina Tune Beads. Sequencing was performed on Illumina NextSeq 2000 instruments with a 2×150 read format with on-board denaturation and targeting 1 million reads per sample.

PhiX (Illumina, catalog #FC-110-3001) was added to each sequencing run as an internal control, aiming for 2-10% alignment. Two positive controls were used each for RSV-A and RSV-B libraries. RSV-A controls included extracted RNA from culture isolate strain A2001-3/12 and ATCC whole genome RNA (catalog #VR-1540DQ). RSV-B controls included extracted RNA from culture isolate strain B1 and ATCC whole genome RNA (catalog #VR-955D). Negative (non-template) controls were nuclease-free water added during extraction. Positive and negative controls were included on each sequencing run and libraries were only sequenced if positive controls showed a visible amplification band, and negative controls had no discernable banding in the 250-750bp region when visualized using D1000 Tapestation (Agilent, catalog #5067-5582) or 1.2% agarose FlashGel (Lonza, catalog #57023). Sequencing runs with >50 % reads passing filter and >50% of bases exceeding Phred quality scores of 30 were accepted for demultiplexing and analysis. These relatively low run-level metrics were selected in order to accommodate potentially low complexity libraries that tend to lower passing filter rates. Stringent sample-level metrics described below indicate the high quality of accepted genomes. Positive controls passed QC if they met the same genome acceptability criteria described below.

### Sanger sequencing of fusion gene

For Sanger sequencing of the fusion gene, 4 primers sets were designed (Table S3). One primer set amplified the entire ∼1650 nucleotide length of the fusion gene coding region. The remaining 3 primer sets were designed to provide 2x tiling of the F gene with amplicon lengths of ∼500 nucleotides. Amplification master mix of Superscipt III Platinum One-Step qRT-PCR Kit reagents (ThermoFisher, catalog #11732020) for F gene amplification consisted of 12.5 μL of 2x Superscript III One-Step PCR reaction mix, 1 μL of 25x Superscript III Platinum enzyme mix, 1.5 μL of 10 µM primer mix containing the forward and reverse primers, 9.5 μL of nuclease-free water, and 0.5 μL of RNA. The fusion gene primers, RSVA-F Fwd2 and RSVA-F Rev1 (Table S3), amplified the region under the following conditions: 55°C for 30 minutes, 94°C for 2 minutes, 40 cycles of 94°C for 30 seconds, 55°C for 30 seconds, and 68°C for 2 minutes, with a final extension at 72°C for 5 minutes. The lid temperature was 105°C and the reaction volume was set to 25 µL. Following amplification, PCR products were purified using the QIAquick PCR Purification Kit (Qiagen, catalog #28106), eluted into 50 µL Qiagen buffer EB and diluted to 4 ng/µL with nuclease-free water. 10 μL of diluted PCR product and 5 μL of a single 5 µM primer was pooled and submitted to Genewiz for Sanger sequencing. Sequence analysis was performed using Geneious Prime 2023.1. Sanger reads were trimmed to remove all ambiguous bases at the ends of the reads and then mapped to the corresponding consensus sequence recovered by the UW-ARTIC panel. A minimum coverage threshold of two Sanger reads per base across the entire F gene was set and a consensus sequence was generated. The resulting Sanger consensus sequence was aligned against the UW-ARTIC consensus using MAFFT v7.490 and pairwise identity was determined.

### Shotgun metagenomic and hybrid capture sequencing for RSV-A and RSV-B

Two different targeted enrichment methods were tested: Illumina Respiratory Virus Enrichment Kit with Respiratory Virus Oligo Panel v2 (Illumina catalog #20100469) and xHYB Respiratory Virus kit (Qiagen catalog #334525). For samples tested with the Illumina kit, extracted RNA was subjected to metagenomic shotgun sequencing using double-stranded cDNA synthesis followed by Illumina DNAPrep tagmentation and 14 cycles of dual-indexed PCR. For samples that failed to generate high quality consensus genomes by shotgun sequencing, targeted enrichment was performed by probe-capture using the Respiratory Virus Oligo Panel (Illumina). For samples tested using the Qiagen panel, cDNA synthesis and library preparation were performed with the Qiaseq xHYB respiratory virus kit according to manufacturer’s specifications with 14 cycles of dual-indexed PCR. Following library preparation, enrichment was performed using xHYB reagents and an RSV-specific probe-capture panel (Qiagen catalog #334575 GeneGlobe ID MXMC-05755Z-0). Finished libraries were sequenced on Illumina NextSeq 2000 instruments with 2×150 reads. Consensus sequences were generated using a custom pipeline that performs pan-respiratory virus reference-based assembly for metagenomic and capture libraries: https://github.com/greninger-lab/revica.

### Genome assembly and bioinformatic analysis

Raw reads were assembled into consensus genomes using a custom bioinformatic pipeline (https://github.com/greninger-lab/rsv_ampseq). Raw reads underwent adapter and quality trimming using fastp v0.23.2. Trimmed reads were then aligned to a multi-sequence fasta file containing references for both RSV-A (GISAID accession EPI_ISL_1647421) and RSV-B (GISAID accession EPI_ISL_1647600) using BBMap v39.01. Subsequently, the trimmed reads were remapped to either the RSV-A or the RSV-B reference, based on whichever exhibited more mapped reads from the initial mapping. Amplicon primers were trimmed from the resulting BAM file using iVar trim v1.4. Variant calling was performed using iVar v1.4 and bcftools v1.17 at a minimum threshold of 0.01, minimum base quality of 20, and minimum depth of 10. A consensus sequence was generated using iVar consensus v1.4 at a minimum frequency threshold of 0.6, a minimum base quality of 20, and a minimum depth of 10, below which ‘N’ bases were called. Maximum likelihood phylogenetic trees were constructed using Nextstrain’s Augur pipeline (v7.2.0) [12], and visualized using Nextstrain’s Auspice tool (<u>v2.61.2</u>) and ggtree (v3.14.0) [13]. Contextual sequences were included by downloading all available RSV-A and RSV-B sequences from NCBI Genbank as of December 19^th^, 2024 and subsampling to include 5 sequences per country per year-month. Lineage assignment was performed using Nextclade (web version 3.10.0) [14,15].

### Genome acceptability criteria

The following criteria were established to determine a passing result from the UW-ARTIC RSV assay. In addition to run-level metrics defined above, to pass QC, a sample must have received at least 1 million raw reads, with at least 500X average per-base depth of coverage across the genome, at least 1000X average per-base depth of coverage across the fusion (F) gene, and >90% of the F gene with over 100X per-base depth of coverage. Consensus genomes generated must have <15% ambiguous bases (Ns).

## Results

### UW-ARTIC amplicon panel for whole genome sequencing of RSV-A and RSV-B

The amplicon panel used in this study was adapted from the ARTIC RSV-A and RSV-B primer schemes described previously [7,8]. Modifications were made to accommodate circulating diversity of RSV using recent samples from Washington state to prevent amplicon dropout (see Methods). The resulting primer scheme (UW-ARTIC) consists of two sets of 50 primer pairs for RSV-A and RSV-B respectively, each set consisting of two pools, and generating fragments ∼400bp in length (Table S1).

### Sensitive recovery of whole RSV-A and RSV-B genomes using the UW-ARTIC Amplicon Panel

To determine the limit of detection for the UW-ARTIC RSV amplicon panel, serial dilutions were prepared using extracted RNA from clinical and cultured samples, and concentrations verified by RT-qPCR (Table 1). For RSV-A, serial dilutions spanned a Ct range of 29-33 (54,394 copies/mL to 238 copies/mL). For RSV-B, Ct values ranged between 27-32 (218,120 copies/mL to 9,544 copies/mL). Multiple replicates (n = 4 to 20, Table 1) were prepared for each serial dilution, and the limit of detection was defined as the concentration at which >95% of replicates generated a genome meeting defined acceptability criteria. Based on this, the limit of detection for both RSV-A and RSV-B was at Ct 30 (487 copies/reaction for RSV-A, 1703 copies/reaction for RSV-B). At Ct 31 (173 copies/reaction for RSV-A, 615 copies/reaction for RSV-B), 6/20 RSV-A and 4/20 RSV-B replicates failed a single metric (<90% of F gene at 100X depth of coverage, Figure S1) while meeting all other genome acceptability criteria.

**Table 1.** Analytical sensitivity of UW-ARTIC RSV amplicon panel.

| **Virus** | **Target Ct** | **Actual Ct\*** | **Viral copies/**<br>**reaction\*** | **Number of**<br>**replicates** | **Passing**<br>**Replicates (%)** |
| --- | --- | --- | --- | --- | --- |
| RSV-A | 29 | 28.55 | 1849 | 4 | 100% |
| RSV-A | 30 | 30.46 | 487 | 20 | 95% |
| RSV-A | 31 | 31.95 | 173 | 20 | 70% |
| RSV-A | 32 | 34.31 | 34 | 20 | 35% |
| RSV-A | 33 | 36.34 | 8 | 5 | 20% |
| RSV-B | 27 | 26.99 | 9270 | 20 | 95% |
| RSV-B | 28 | 27.92 | 4796 | 20 | 100% |
| RSV-B | 29 | 28.95 | 2873 | 20 | 100% |
| RSV-B | 30 | 29.83 | 1703 | 20 | 100% |
| RSV-B | 31 | 30.94 | 615 | 20 | 80% |
| RSV-B | 32 | 32.14 | 406 | 5 | 0% |

\*As measured by quantitative real-time RT-PCR of the matrix gene

### High specificity and no cross reactivity to non-RSV respiratory viruses

The analytical specificity of the UW-ARTIC-RSV Amplicon panel was determined using clinical samples that tested negative for both RSV-A and RSV-B (n=20), and a set of non-RSV respiratory virus samples (n=20, Table 2). This included clinical remnant samples that were positive for seasonal coronaviruses, rhinovirus, human parainfluenza virus 1, human parainfluenza virus 3, human parainfluenza virus 4, SARS-CoV-2, adenovirus, metapneumovirus, influenza A virus, and influenza B virus. Nonspecific banding was observed in several of the non-RSV samples but no RSV-A or RSV-B genomes were recovered (Table 2), with the number of reads mapping to the genome equaling <0.5% of trimmed reads and covering <1.5% of the genome in all cases (Table S4), with most mapped reads representing primer artifacts. No RSV genomes were recovered from clinically negative samples (Table 2 and S4).

**Table 2.** Specificity of UW-ARTIC RSV amplicon panel. No recovered genomes from RSV-negative, and non-RSV positive specimens.

| Virus | n | Recovered genomes |  |
| --- | --- | --- | --- |
|  |  | RSV-A | RSV-B |
| RSV-negative samples | 20 | 0 | 0 |
| Coronavirus (non-SARS-CoV-2) | 2 | 0 | 0 |
| Rhinovirus | 2 | 0 | 0 |
| Parainfluenza virus 1 | 1 | 0 | 0 |
| Parainfluenza virus 3 | 2 | 0 | 0 |
| Parainfluenza virus 4 | 1 | 0 | 0 |
| SARS-CoV-2 | 5 | 0 | 0 |
| Adenovirus | 2 | 0 | 0 |
| Metapneumovirus | 2 | 0 | 0 |
| Influenza A | 2 | 0 | 0 |
| Influenza B | 1 | 0 | 0 |
| TOTAL | 40 | 0 | 0 |

### Robust performance with clinical samples across a range of viral loads

To assess the performance and breadth of genome recovery of this panel, 20 RSV-A and 20 RSV-B positive clinical samples were randomly selected. These were remnants of clinical swab specimens that were collected between October 2022 to February 2023 and stored at −80°C. Samples had viral loads between 3.98 × 10^3^-6.75 × 10^7^ copies/mL (Table S5), i.e. 6 samples (2 RSV-A, 4 RSV-B, Table S5) had viral loads below the limit of detection of this assay.

Of the 40 sequenced samples, 19 RSV-A (95%) and 17 RSV-B samples (85%) generated genomes meeting all defined acceptability criteria. Due to the split pool design, there was variation in read depth across the genome (Figure S2), however mean depth of coverage from both pools was sufficient to meet required thresholds for mean coverage in all cases (>500X across the genome and >1000X in F, Figure 1). Across all 40 samples, the percentage of mapped reads ranged from 50.3 to 93.6% (median 81.0%) and over 92% of the genome had at least one read mapping to the reference (median 99.1%, range 92.8 – 99.4%). Among samples that met acceptability criteria, this resulted in consensus genomes with <3% Ns (median 0.9%, range 0.9 – 2.7%) where Ns were called when there were fewer than 10 reads of support.

**Figure 1.**
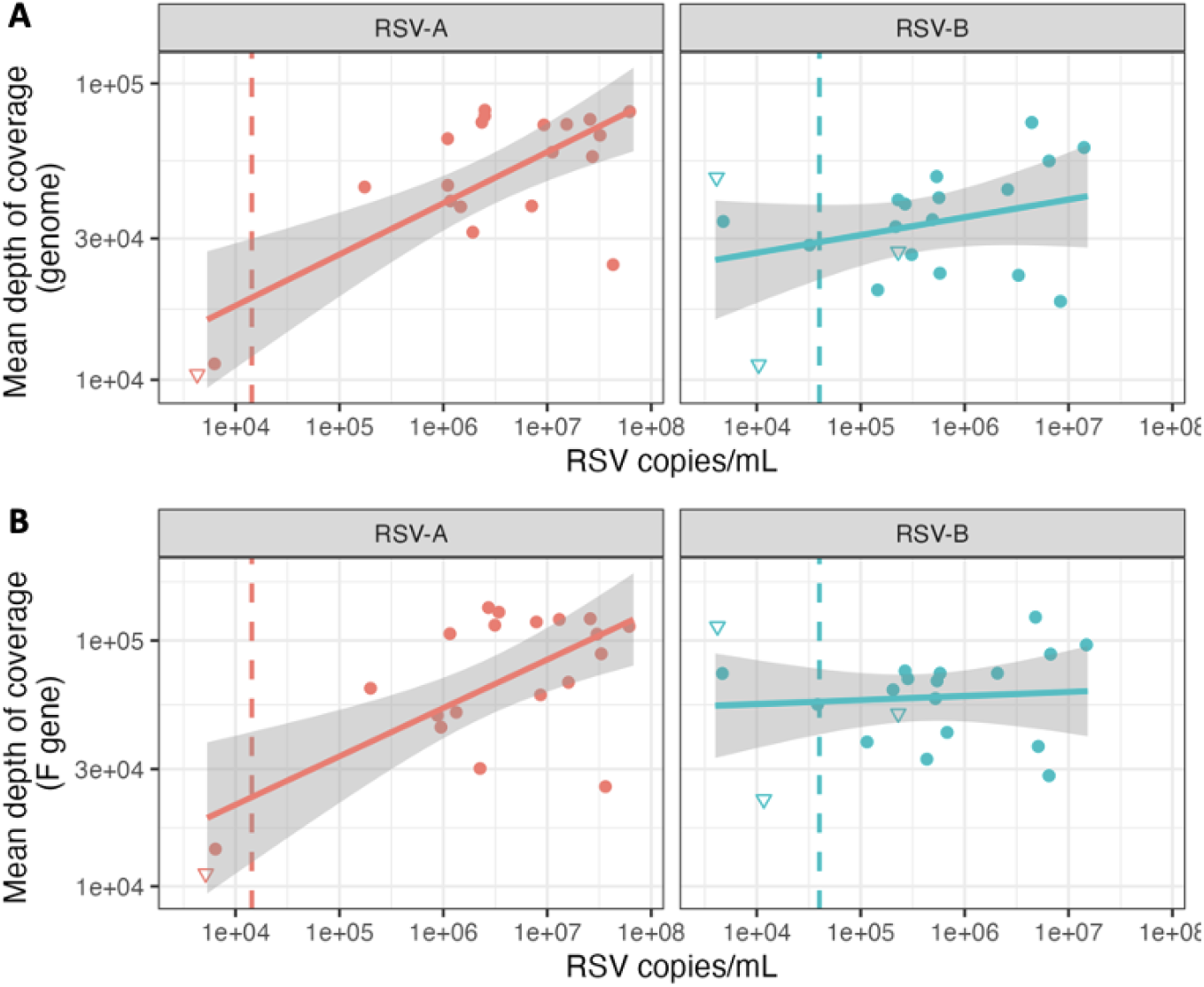
High mean depth of coverage achieved with the amplicon panel both across the genome (A) and within the F gene (B). Dashed line represents limit of detection from sensitivity analysis (Table 1), triangles represent samples that failed genome acceptability criteria.

Among samples that failed to meet passing criteria, all had viral copies below the limit of detection. While each of the four samples that failed QC failed to have at least 100X average depth of coverage at >90% of positions in the fusion gene, 3 out of 4 samples generated a consensus sequence that was adequate for surveillance and lineage identification purposes with <10% ambiguous bases (Ns) in the consensus genome, and complete sequence recovery in the F gene.

Sequenced samples were representative of regional circulating genetic diversity. Nextclade designations for passing consensus genomes included six RSV-A lineages: A.D.1.4 (n=1), A.D.3 (n=4), A.D.3.1 (n=1), A.D.5 (n=1), A.D.5.2 (n=12), and one RSV-B lineage B.D.E.1 (n=20) (Table S5). Validation samples clustered with other sequences from Washington State (Figure 2).

**Figure 2.**
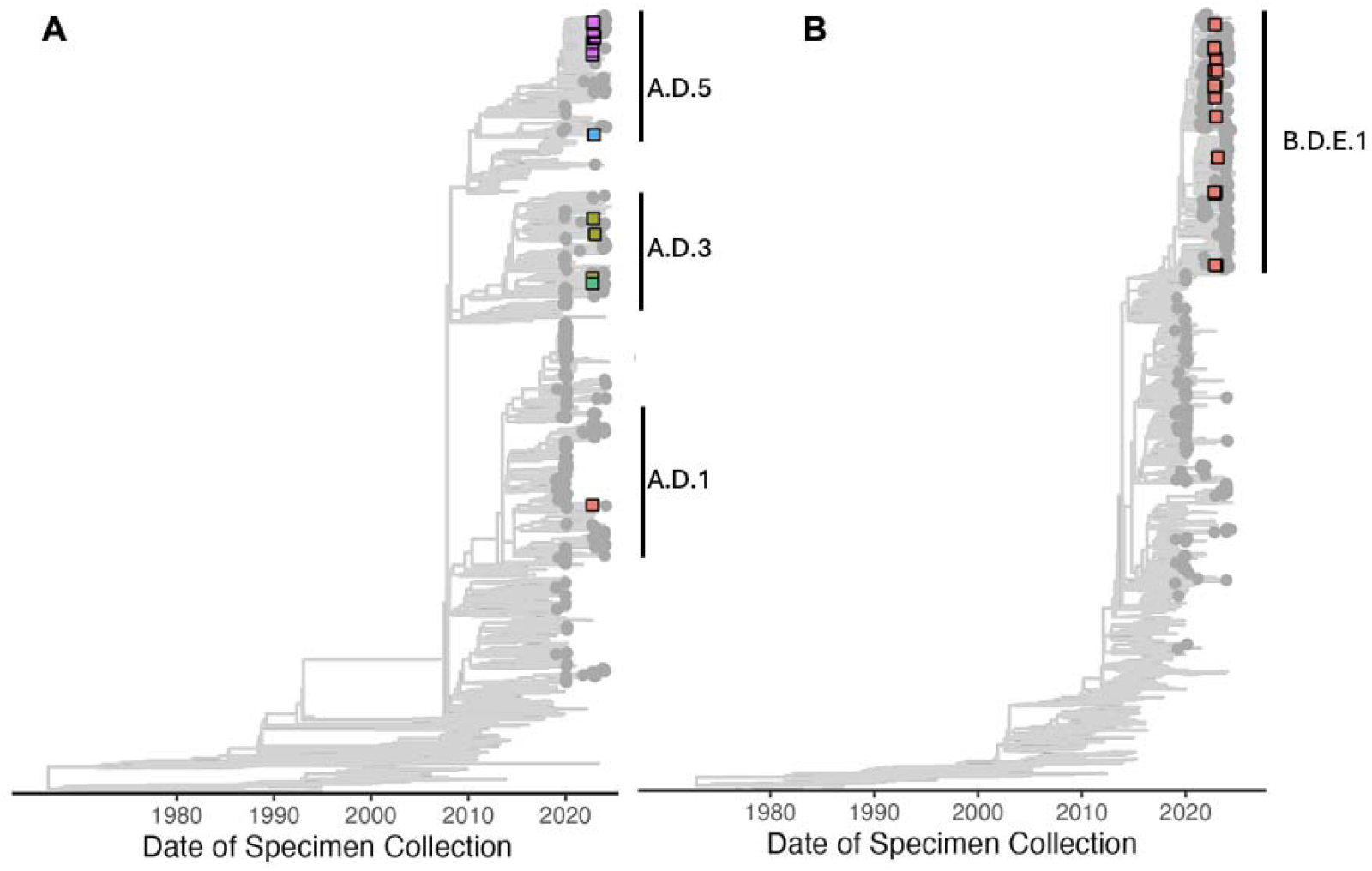
Maximum likelihood trees of RSV-A (A) and RSV-B (B). Square tip shapes indicate study samples (n=19 RSV-A; 20 RSV-B), grey tip shapes represent other publicly available sequences from Washington, Oregon, and Idaho (n=608 RSV-A; 592 RSV-B); and other global sequences are shown without tip shape (n = 1678 RSV-A; 1596 RSV-B).

### Amplicon dropouts in some samples potentially due to mutations in primer binding regions

Mutations occurring in primer regions can affect PCR amplification leading to amplicon dropout or reduced coverage, and this can impact genome completeness. Potential amplicon dropout was observed in 5 out of 50 primer pairs for RSV-A; and 10 primer pairs for RSV-B affecting 1 to 5 samples in each case (Table S6). Dropouts were identified by looking for strings of Ns in the consensus sequence generated when there are fewer than 10 reads of support for a given position. A full amplicon dropout was defined as contiguous strings of Ns exceeding 200bp within the amplicon. The largest number of dropouts were seen in RSV-A amplicon 47 (n=5) and RSV-B amplicon 29 (n=3) both in the L gene. Four out of 5 samples with dropout of RSV-A insert 47 and 1 out of 3 samples with dropout of RSV-B insert 29 had a mutation near the 31 end of one of the primers (Figure S3 and S4). Mutations in primer binding regions were therefore not the sole reason for amplicon dropout. Samples with dropouts generally had lower viral copies compared to those with complete recovery of the amplicon (average 1.87 × 10^7^ copies/mL for complete RSV-A insert 47 vs. 2.23 × 10^6^ copies/mL for dropout; 2.76 × 10^6^ copies/mL for complete RSV-B insert 29 vs. 2.02 × 10^5^ copies/mL for dropout). No dropouts were observed within the F gene for RSV-A. For RSV-B, a single sample, RSV-B 14, had a dropout for amplicon 19, but also had other low coverage regions likely due to low viral load (5.10 × 10^3^ copies/mL).

### Sensitive detection of intra-host variation

To interrogate the limit of detection of minor alleles in mixed populations, two samples were selected that had similar viral loads and at least 8 single nucleotide variants (SNVs, including polymorphisms, insertions and deletions) distinguishing them (Table 3). Extracted RNA from the two samples at similar viral loads (within 0.3 Ct difference) was mixed at varying volume ratios: 0:100, 1:99, 5:95, 10:90, 25:75, and 100:0 of sample 1 to sample 2 (Table 4). Mixed samples were run in triplicate and results were reviewed to determine whether the distinguishing SNVs were identified at greater than 1% allele frequency.

**Table 3.** Expected SNVs in RSV-A samples (A1 and A2) and RSV-B samples (B1 and B2)

| Sample | SNVs |
| --- | --- |
| **A1** | C5747T, C5765T, G5988A, A6086G, A6416T, A6701G |
| **A2** | C5691T, C5274T, C5900T, G6080A, C6794T, T6872A, C6896T, A7076G, T7202C, T7223C, A7349T, T7361C |
| **B1** | T5791C, G5841A, T7243C, C7561T |
| **B2** | A7257G, C7443T, G6448A, T6470C |

**Table 4.** Sample mixtures and recovery of SNVs at allele frequencies of 1% or higher with at least 10 reads of support.

| Subtype | Sample ID | %A1 | %A2 | A1 SNVs detected<br>(6 expected) | A2 SNVs detected<br>(12 expected) |
| --- | --- | --- | --- | --- | --- |
| **RSV-A** | MIX1 | 0 | 100 | 0 | 12 |
| **RSV-A** | MIX2 | 1 | 99 | 6* | 12 |
| **RSV-A** | MIX3 | 5 | 95 | 6 | 12 |
| **RSV-A** | MIX4 | 10 | 90 | 6 | 12 |
| **RSV-A** | MIX5 | 100 | 0 | 6 | 0 |
| Subtype | Sample ID | %B1 | %B2 | B1 SNVs detected<br>(4 expected) | B2 SNVs detected<br>(4 expected) |
| **RSV-B** | MIX1 | 0 | 100 | 0 | 4 |
| **RSV-B** | MIX2 | 1 | 99 | 1** | 4 |
| **RSV-B** | MIX3 | 5 | 95 | 4*** | 4 |
| **RSV-B** | MIX4 | 10 | 90 | 4 | 4 |
| **RSV-B** | MIX5 | 25 | 75 | 4 | 4 |
| **RSV-B** | MIX6 | 100 | 0 | 4 | 0 |

\*G5988A detected at allele frequencies lower than 1% in all 3 replicates; \*\*T7243C detected in 2 out of 3 replicates, all others were either detected at frequencies lower than 1%, or with fewer than 10 reads of support; \*\*\*C7561T detected in 1 out of 3 replicates; G6448A and T6470C detected in 2 out of 3 replicates

Using an allele frequency cutoff of 1%, all expected SNVs were identified at mixing ratios of 5:95 and higher for both RSV-A and RSV-B (Table 4, Figure 3). At mixing ratios of 1:99, 5 out of 6 expected RSV-A SNVs were identified—a single variant (G5988A) was detected at allele frequencies lower than 1%, which is expected based on the stochasticity of mixing at 1% allele frequency. For RSV-B, at mixing ratios of 1:99, most variants were detected at allele frequencies lower than 1%, or with fewer than 10 reads of support, which is expected due on the stochasticity of mixing at 1% allele frequency and one variant (T7243C) was detected in 2 out of 3 replicates.

**Figure 3.**
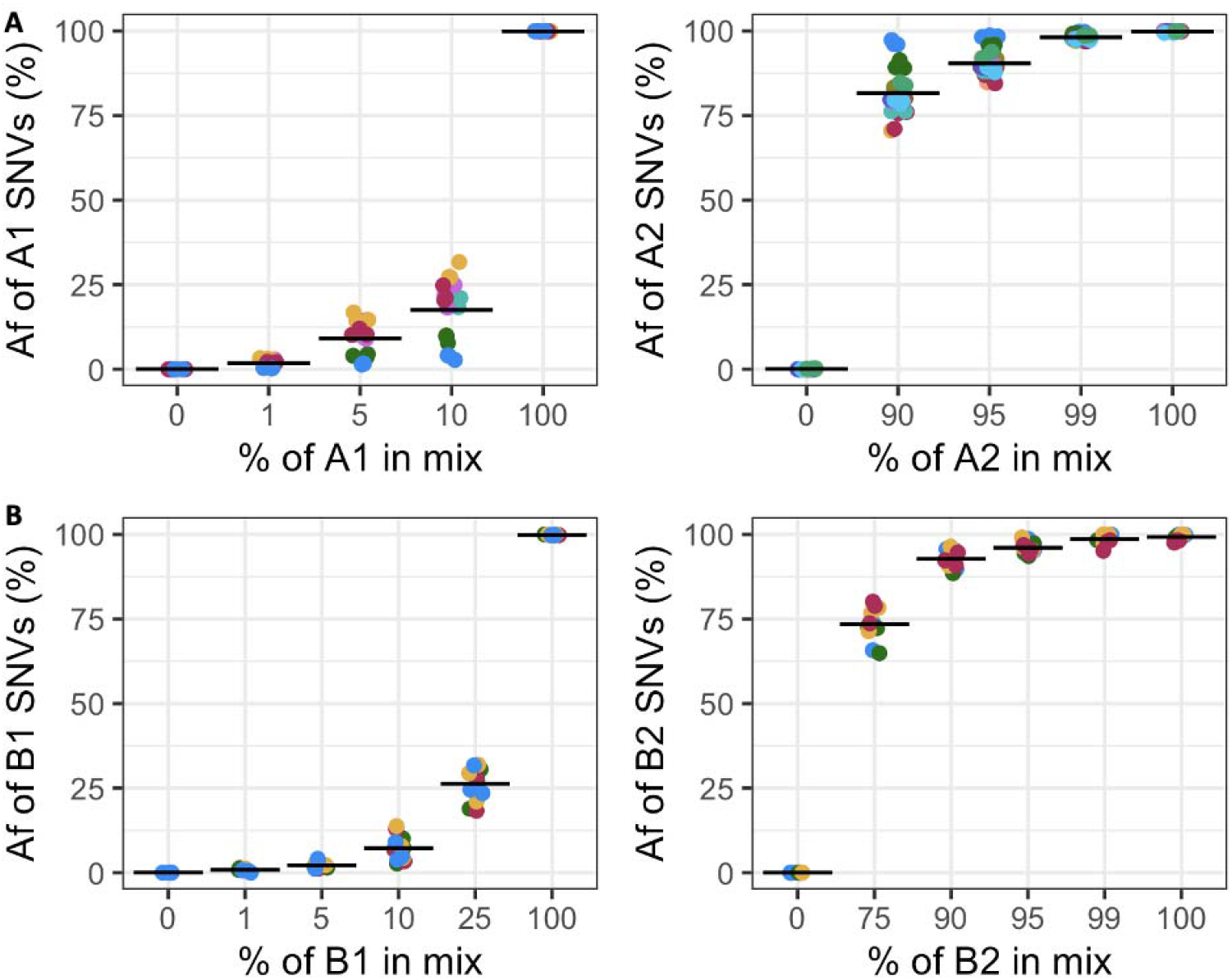
Detection of intra-host SNVs defining selected samples (Table 3) for mixtures of RSV-A (A) and RSV-B (B) as defined in Table 4. Dot colors within each panel represent different SNVs. Af = allele frequency.

### High inter-run and intra-run assay precision

Assay precision was determined by preparing and sequencing clinical samples in duplicate, either by the same technician on the same run (repeatability or intra-run precision), or by different technicians on separate runs (reproducibility or inter-run precision). Performance was determined by comparing consensus genomes and allele frequencies of variant calls for all sample pairs that met genome acceptability criteria.

For all sample pairs meeting genome acceptability criteria (repeatability: n=18 pairs for RSV-A, n=8 for RSV-B; reproducibility: n=15 pairs for RSV-A, n=7 for RSV-B) repeated samples were highly concordant at both the consensus (Table S7 and S8) and sub-consensus level (Figure 4). Consensus genomes had 100% pairwise identity with 0-2 pairwise differences (median 0 differences) across the whole genome after masking regions of low coverage that were called as Ns (Table S7 and S8).

**Figure 4.**
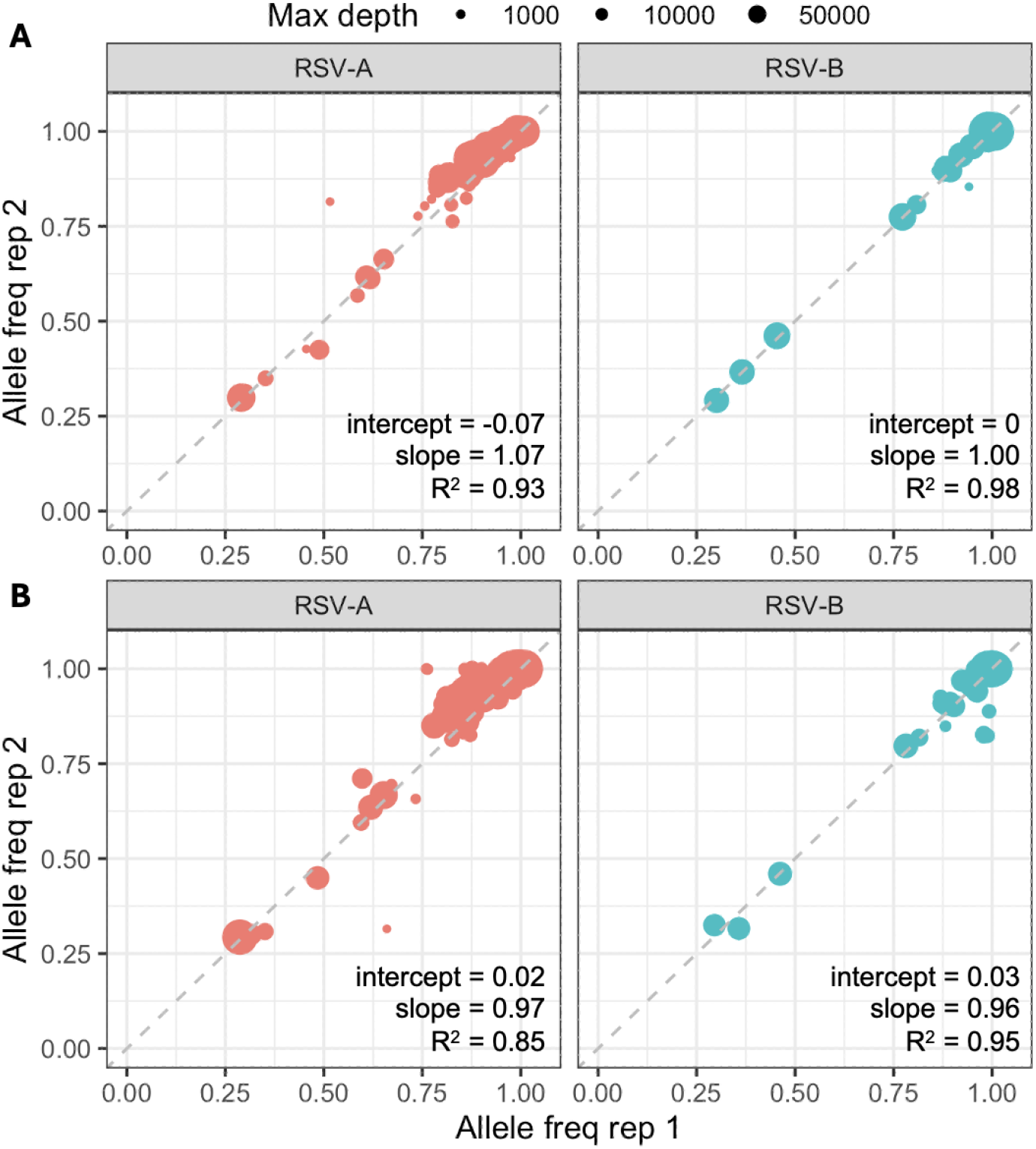
Assay precision determined by comparing allele frequency of variants identified in repeated samples either on the same run A) repeatability (intra-run precision) or different runs B) reproducibility (inter-run precision). Each point indicates a single mutation, size of circles indicates maximum depth of coverage at a given SNV position across the two replicates. Figure shows all mutations called at 1% allele frequency or higher in all sample pairs.

Comparing variants called at 1% frequency or higher and excluding variant calls with fewer than 10 reads of support or fewer than 100 total reads covering the position, there was a high level of concordance between allele frequency of variants identified between replicates (Figure 4). Combining variants called in all sample pairs, a mean of 97.0% (n = 24) and 97.5% (n=20) of variants were called across replicates intra-run and inter-run respectively. The allele frequencies of the variants called were highly concordant between replicates, with R^2^ values ranging between 0.85 and 0.98 (Figure 4)

### Accurate recovery of F gene sequence compared to alternative reference sequencing methods

To determine the accuracy of the UW-ARTIC amplicon sequencing approach, results were compared to existing methods including shotgun metagenomic sequencing, hybridization capture sequencing, and Sanger sequencing of the F gene, due to the relevance of the fusion protein as an antiviral and vaccine target.

Shotgun metagenomic sequencing was attempted on 6 RSV-A samples and 6 RSV-B samples with Ct <25. All 6 RSV-A samples produced sequences that met genome acceptability criteria, while only 2 RSV-B samples produced sequences that met genome acceptability criteria. All samples that met these sequencing requirements had F gene sequences that were identical to the sequence recovered by amplicon sequencing (Table 5).

Two different hybridization capture panels were tested for RSV whole genome recovery. After down-sampling to the same number of raw reads, the Illumina and Qiagen panels produced similar depth of coverage across the genome (Figure S5). The Illumina panel was used to perform hybridization capture sequencing on 7 RSV-A samples and 18 RSV-B samples, including the 4 RSV-B samples that failed to meet genome acceptability criteria from shotgun sequencing. Across all capture samples, the F gene sequence was identical to the corresponding sequence recovered by amplicon sequencing (Table 5).

Sanger sequencing was performed on 15 RSV-A samples and the complete sequence of the F gene was recovered. Sanger sequencing of the F gene was attempted for RSV-B samples but was unsuccessful due to little to no amplification in attempted samples. The F gene sequence was identical between sanger and amplicon sequencing for 14 out of 15 RSV-A samples (Table 5). In the discrepant sample, a mixed base (A/T) was called in the Sanger sequence with 2 supporting reads while the amplicon NGS call was primarily A with other minor alleles at < 1% at that position (allele frequencies A: 99.5% C: 0.2% T: 0.1% G: 0.2%; reads: 30204, 68, 24, 48; and 6 reads with Ns). Capture and shotgun sequencing results on the discrepant position supported the amplicon NGS call.

Overall, the F gene sequences from UW-ARTIC amplicon sequencing had > 99% pairwise identity to sequences obtained by alternative methods (Table 5).

**Table 5:**
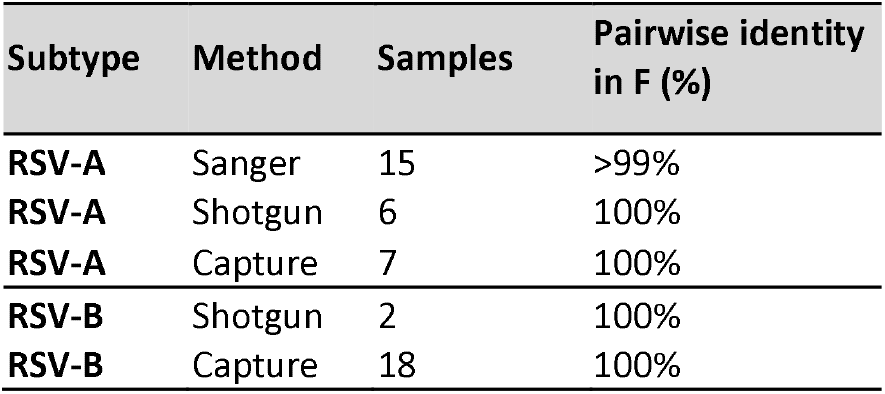
Comparing F gene consensus sequences obtained by alternative methods.

## Discussion

Using guidelines from the US Food and Drug Administration, Clinical Laboratory Improvement Amendments, and College of American Pathologists for validating assays for use in clinical settings, we have determined the performance characteristics of the UW-ARTIC-RSV amplicon panel to produce high-quality genomes. The panel is a modified version of the ARTIC RSV panel [7,8] with changes to primers based on circulating RSV diversity primarily in Washington State, making it more robust against US strains of RSV. However, these assay updates are likely to improve performance against globally circulating strains of RSV as well. For example, in an alignment of 2370 global RSV-B sequences from 2021-2024, the updates to RSV-B primers 17 and 45 represent mutations in a majority of recent sequences: 4948G, 5336A, and 13734T are present in 85.7%, 87.2% and 97.2% sequences respectively. We have also developed a robust bioinformatic pipeline for genome assembly, quality control, and variant calling. Although other whole genome sequencing methods exist for RSV [5,16], this is, to our knowledge, the first clinically validated RSV sequencing assay with design characteristics allowing for sensitive genome recovery. It has been used to generate FDA-reportable data in clinical trials for RSV antiviral products. The assay can recover high-quality genomes from low viral load samples, with a limit of detection that is similar to existing amplicon panels [16,17]. By leveraging Illumina COVIDseq/MAP kits and automated robotic liquid handling systems, assay throughput can be rapidly scaled up for large volume or surge testing.

The UW-ARTIC RSV panel shows high precision and accuracy with the ability to consistently generate sequences that have high pairwise identity in repeated sequencing attempts, accurate recovery of the F gene compared to alternative methods, and sensitive recovery of minor variants. High quality consensus genomes are important for genomic surveillance, contact tracing, and identifying mutations of concern. In addition, the ability to accurately and sensitively identify minor variants in allelic mixtures is important for tracking treatment emergent mutations in clinical trials of vaccines and antiviral drugs, and in tracking viral evolution during prolonged infection, such in immunocompromised individuals. Alternative approaches such as shotgun and probe-capture based sequencing offer advantages like being able to recover a broader range of targets, but they come with higher costs, longer workflows, and lower analytical sensitivity, requiring higher viral load samples and more reads per sample to achieve comparable depth of coverage for clinical applications.

A limitation of the assay is the use of separate primer sets for RSV-A and RSV-B, necessitating RSV typing [18] prior to library preparation, however work with multiplexing primer sets has shown positive results so far [7] and future designs may include both RSV-A and RSV-B in a single primer set. Secondly the split pool design has the potential to produce uneven coverage across the genome, which can be mitigated by adjusting concentrations of the two pools, performing additional primer balancing and quality control, and quantifying PCR products prior to combining pools to ensure equimolar pools. Single-tube [9] and cross-subtype designs would improve the efficiency of this workflow but present potential challenges in PCR optimization. Thirdly, we did not compare results from the UW-ARTIC RSV panel against other available amplicon schemes ([16,17]). Finally, like other amplicon designs, our results show that mutations in primer binding regions, viral load, and potentially other factors such as primer pooling issues can impact amplicon dropout. Separate from this study, we have used the UW-ARTIC RSV panel to sequence samples from clinical trial sites around the globe from the 2023/24 respiratory virus season and no widespread dropouts or quality issues were observed. However continuous genomic surveillance to identify mutations in primer binding regions along with regular updates and revalidation of primer pools as well as methods for primer balancing will ensure continued utility and high performance of this and similar amplicon panels.

