## Supplementary Figures for "Clinical performance evaluation of a tiling amplicon panel for whole genome sequencing of respiratory syncytial virus"

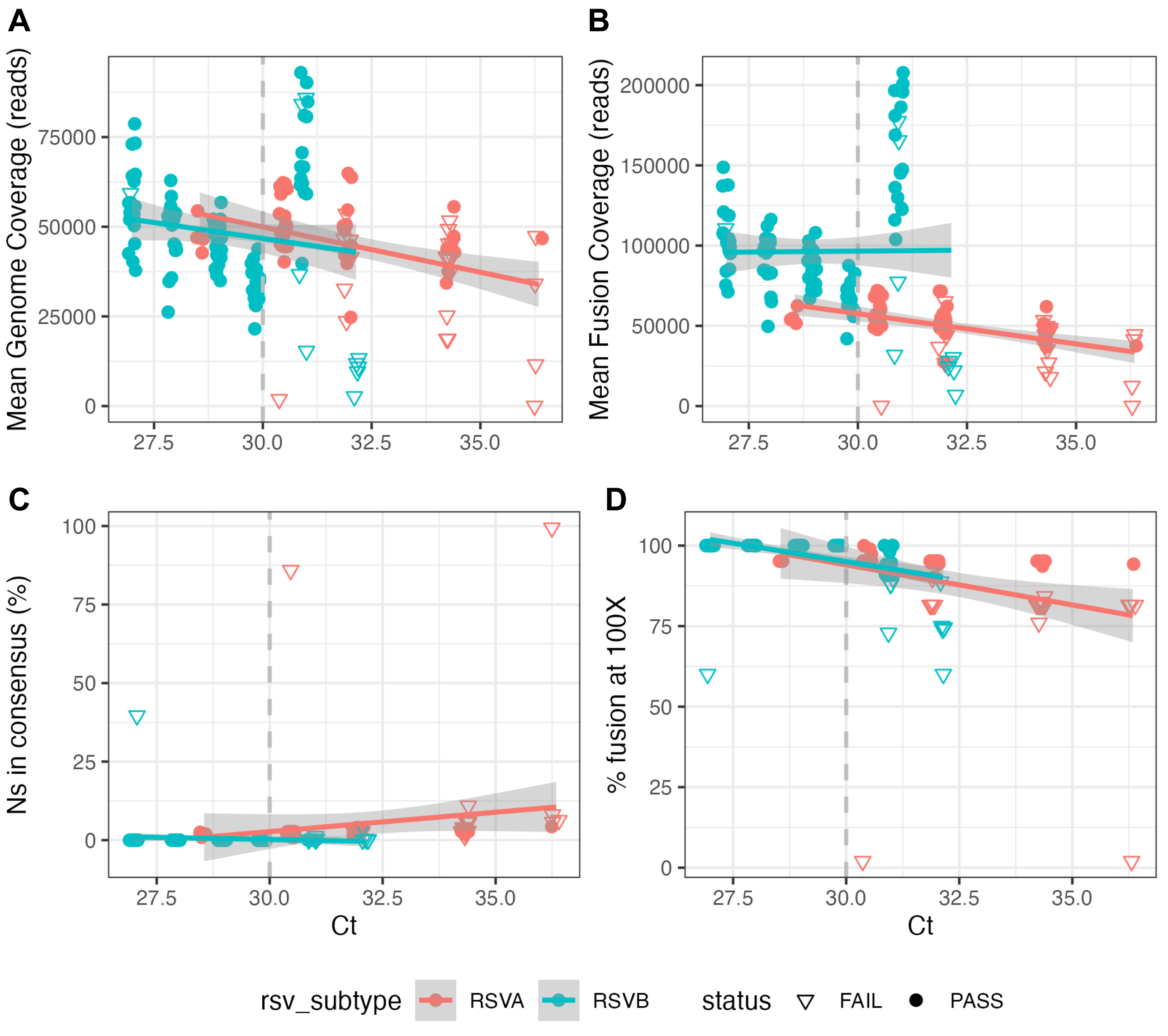


**Figure S1:** UW-ARTIC RSV Amplicon Panel Sensitivity. Mean per-base depth of coverage across the whole genome (A) or F gene (B); (C) genome completeness; (D) % of F gene with >100X depth of coverage. Dashed line indicates limit of detection based on >=95% of replicates meeting all QC criteria.

A

B

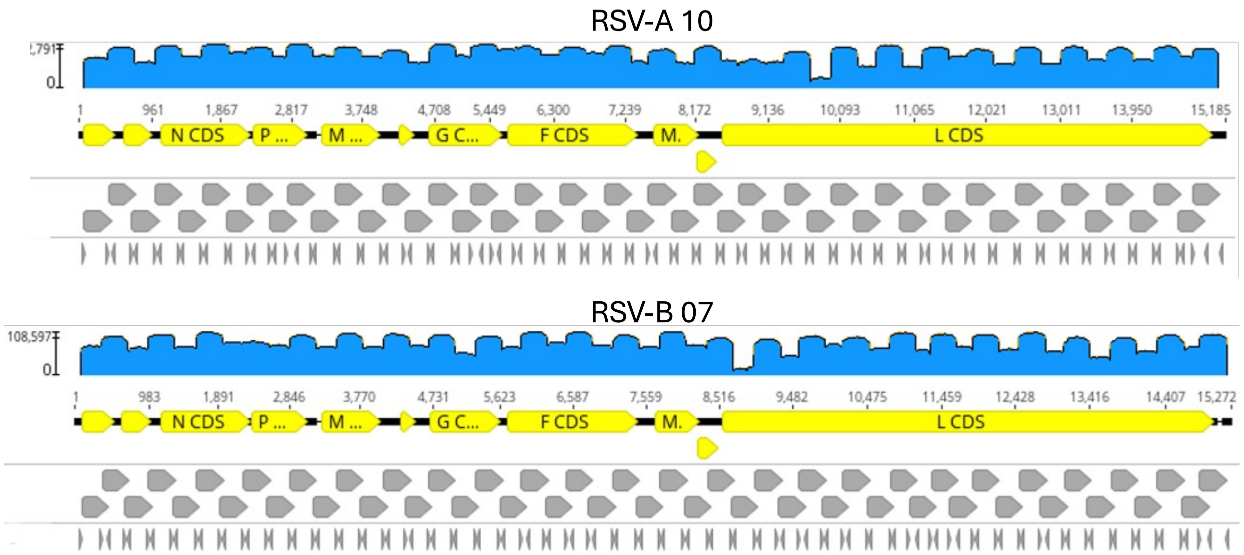


**Figure S2:** Differences in coverage across the genome due to 2-pool design.

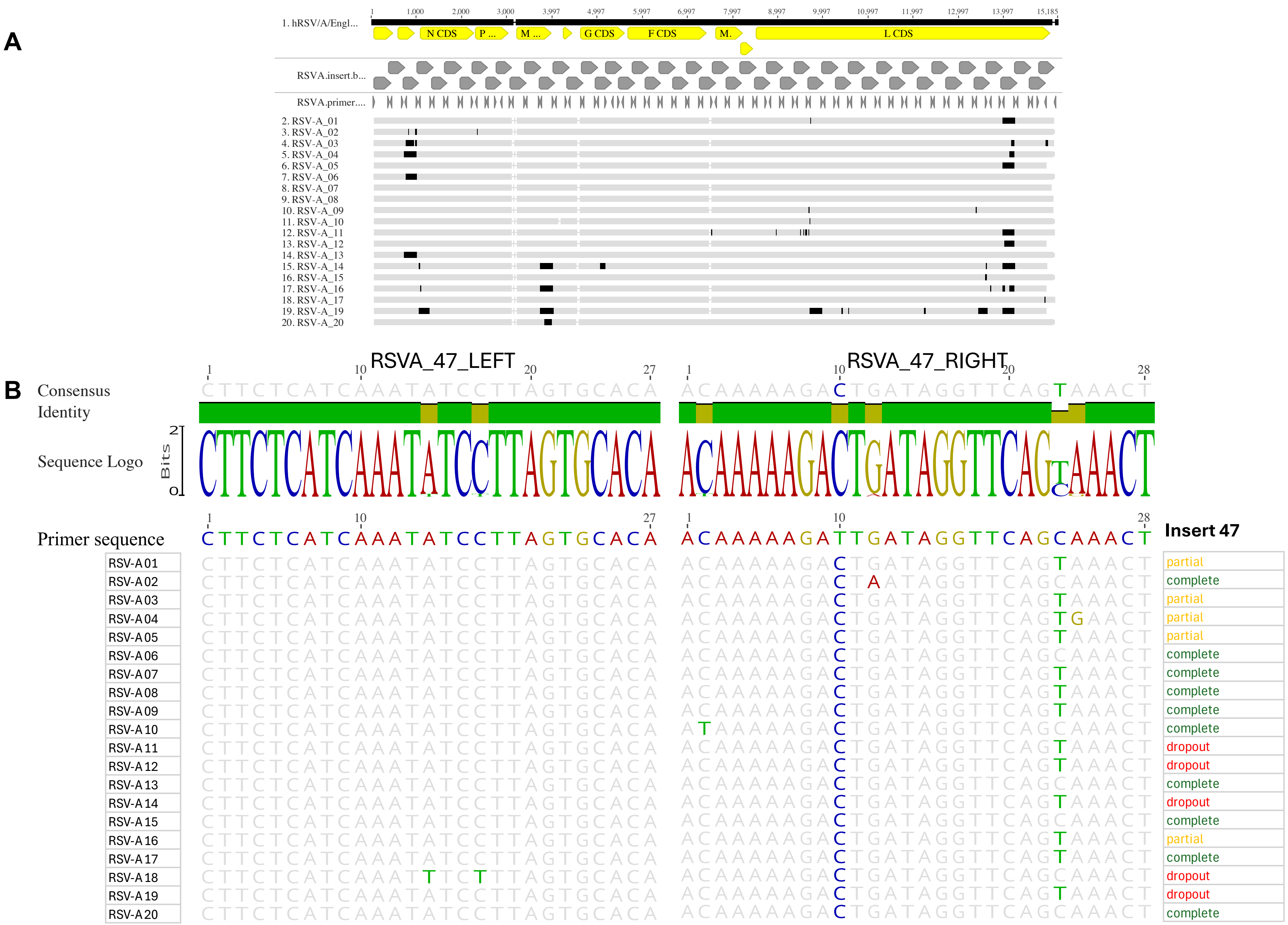


**Figure S3**. (A) Alignment of 19 RSV-A genomes with dropouts indicated in black (B) Variation in primer region where amplicon dropout was observed for RSV-A in insert 47. Primer sequences are shown in 5'-3' orientation.

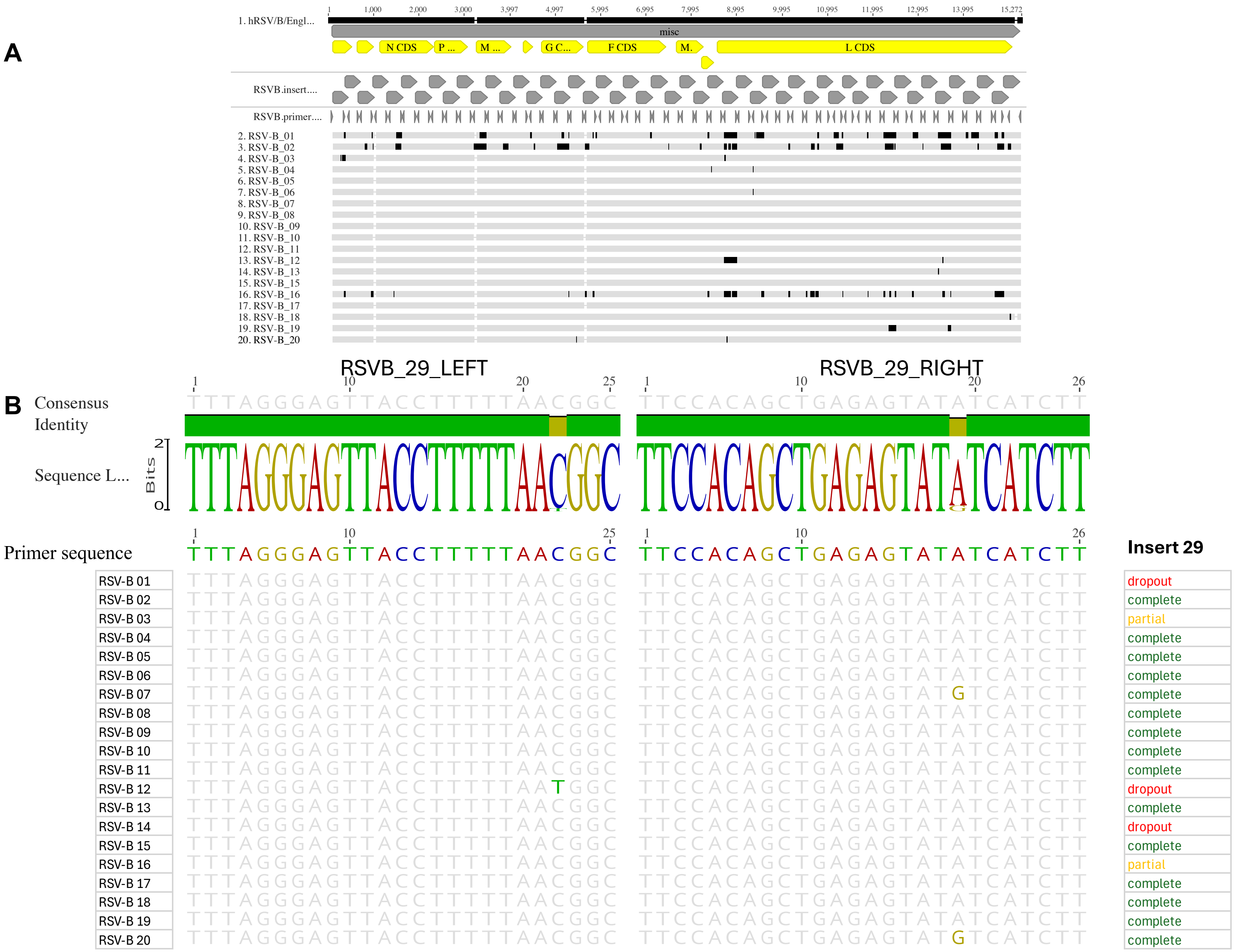


**Figure S4**. (A) Alignment of 19 RSV-B genomes with dropouts indicated in black. (B) Variation in primer region where amplicon dropout was observed for RSV-B in insert 29. Primer sequences are shown in 5’-3’ orientation.

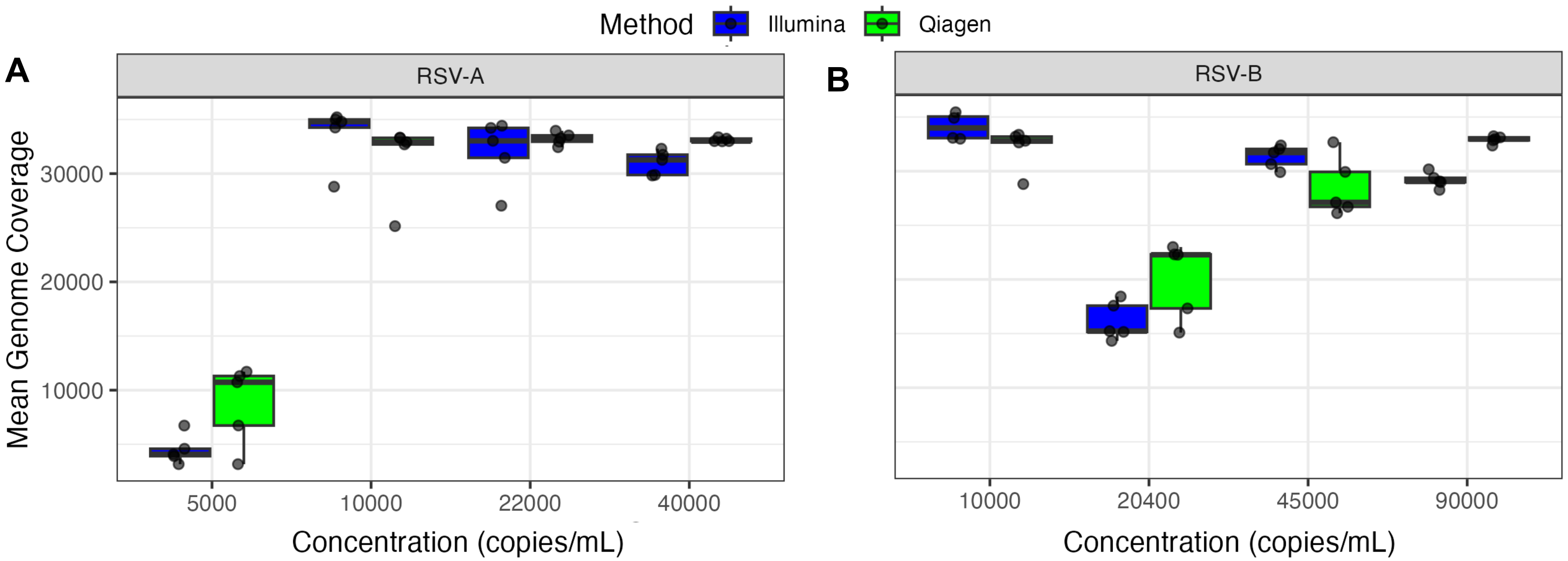


**Figure S5**. Comparing performance between Illumina and Qiagen panels for whole genome capture sequencing of RSV-A (A) and RSV-B (B).
