## Supplementary Tables for "Clinical performance evaluation of a tiling amplicon panel for whole genome sequencing of respiratory syncytial virus"

Table S1. UW-ARTIC RSV amplicon panel

| Pool | RSV A Primer | Primer Sequence | Pool | RSV B Primer | Primer Sequence |
| --- | --- | --- | --- | --- | --- |
| A1 | RSVA_1_LEFT | TTGGTTAGAGATGGGCAGCAAC | B1 | RSVB_1_LEFT | TGATGAGTGCTATTTAAGTCTAACCTTT |
| A1 | RSVA_1_RIGHT | AGGTCAAATCCAAGTAATTCAGATAATTGA | B1 | RSVB_1_RIGHT | AGTCATTACTGAGTCACTTAGTCTTTTAGA |
| A1 | RSVA_3_LEFT | TCTAACCAGAGACATCATAACACATAAATT | B1 | RSVB_3_LEFT | AACGACAACACCACCATGCAAA |
| A1 | RSVA_3_RIGHT | CCCATCTTTCATCTTATGTCTCTCCT | B1 | RSVB_3_RIGHT | TTGGAGAATGGCTTGGTTTGGA |
| A1 | RSVA_5_LEFT | GCTATGTCTAGATTAGGAAGAGAAGACA | B1 | RSVB_5_LEFT | ACACCTAAACAAACTATGTGGTATGC |
| A1 | RSVA_5_RIGHT | TCCTTGGGTAATAAACCTTTATAACGTT | B1 | RSVB_5_RIGHT | ACCTGATCTATCTCCTGCTGCT |
| A1 | RSVA_7_LEFT | ATGGAACAAGTTGTGGAGGTGT | B1 | RSVB_7_LEFT | AGGACAAGTAATGCTAAGATGGGG |
| A1 | RSVA_7_RIGHT | CTTCTCCATGGAATTCAGGAGCA | B1 | RSVB_7_RIGHT | TCTTTGGGGTTGAGTTGATGCT |
| A1 | RSVA_9_LEFT | AGATGATACTGTAGGGAACAAGCC | B1 | RSVB_9_LEFT | CCGATAACATCTGGCACCAACA |
| A1 | RSVA_9_RIGHT | TCATTCCTGAGTCTTGCCATAGC | B1 | RSVB_9_RIGHT | CATTGGTCATTAATGCTTCTGCTCT |
| A1 | RSVA_11_LEFT | ACACAACACCAATAGAAAACCAACA | B1 | RSVB_11_LEFT | CCATTGAATCAATTGCCAGACTGA |
| A1 | RSVA_11_RIGHT | TGCTAGCACTGCACTTCTTGAG | B1 | RSVB_11_RIGHT | TGGTGAAATTACTAGGCATTTGAGC |
| A1 | RSVA_13_LEFT | ACCAACATACCTAAGATCCATCAGC | B1 | RSVB_13_LEFT | ACCAACCTATCTAAGATCAATCAGTGT |
| A1 | RSVA_13_RIGHT | TGAGTTTTGTTTGATTGATTGAACCAC | B1 | RSVB_13_RIGHT | TGAGTGGGATTGGTTTGTGTGA |
| A1 | RSVA_15_LEFT | TCTGGCCTTACTTTACACTAATACACA | B1 | RSVB_15_LEFT | TTGGCCCTATTTTACACTAATACATATGA |
| A1 | RSVA_15_RIGHT | AGATGATTGAGAGTGTCCCAGGT | B1 | RSVB_15_RIGHT | GTATCCCAGGTCTTTTCTAGAGTCC |
| A1 | RSVA_17_LEFT | CTTCTCCAATCTGTCCGGAACT | B1 | RSVB_17_LEFT | GTCTCACCAGAAAGGGTTGGCC |
| A1 | RSVA_17_RIGHT | TGATGGTTGGCTTTCCTGTAGG | B1 | RSVB_17_RIGHT | TTTTTCGGTGTTTTGGCTAGTG |
| A1 | RSVA_19_LEFT | GCTATCCAAGCCCATCACAAGT | B1 | RSVB_19_LEFT | CGAGCCCTCCACATCAAATTCT |
| A1 | RSVA_19_RIGHT | AGTTATAACACTAGTATACCAACCAGTTCT | B1 | RSVB_19_RIGHT | CTGTAATTCAGTTACTGCATTCTTATACTTATC |
| A1 | RSVA_21_LEFT | ACACTCAACAATACCAAAAACACCA | B1 | RSVB_21_LEFT | ACACCTTGAAGGAGAAGTGAACA |
| A1 | RSVA_21_RIGHT | ACATATAAGTGCTTACAGGTGTAGTTAC | B1 | RSVB_21_RIGHT | AGACATGATAGAATAACTTTGTTGCCTT |
| A1 | RSVA_23_LEFT | CAACCAACACAAAGGAAGGATCC | B1 | RSVB_23_LEFT | TCCTTCTTTCCACAAGCTGACA |
| A1 | RSVA_23_RIGHT | TACAGACACAGTATCCACCCCC | B1 | RSVB_23_RIGHT | CATCAGAAGGAAACACTAGAGGGT |
| A1 | RSVA_25_LEFT | TCCACCACAAATATCATGATAACTACCA | B1 | RSVB_25_LEFT | AGCAAAGACCAACTAAGTGGAATCA |
| A1 | RSVA_25_RIGHT | TTGCAAGGATTCCTTCGTGACA | B1 | RSVB_25_RIGHT | CATGAAGTTTTGCCTCACTAGCA |
| A1 | RSVA_27_LEFT | AGAGTATGCCCTCGGTGTAGTT | B1 | RSVB_27_LEFT | ACCCAATTCACCTAAGATAAGAGTGT |
| A1 | RSVA_27_RIGHT | GGAATTTATACTACAAGGATATTTGTCAGGT | B1 | RSVB_27_RIGHT | TGGCATCTAATAAGTTTTTAGGTGTCC |
| A1 | RSVA_29_LEFT | AGTGTCATAATACTCAATCCTAATACTTACCA | B1 | RSVB_29_LEFT | TTTAGGGAGTTACCTTTTTAACGGC |
| A1 | RSVA_29_RIGHT | CTTTTTGAGTAAATTAGTAGTAGTAGTCTGTTC | B1 | RSVB_29_RIGHT | TTCCACAGCTGAGAGTATATCATCTT |
| A1 | RSVA_31_LEFT | GCAGACAAAAATCAATCCACAAAACA | B1 | RSVB_31_LEFT | AGATAATCAAACTTTGAGTGGTTTTCAGT |
| A1 | RSVA_31_RIGHT | ACATTATTGAATCCACATCTTAAGCCT | B1 | RSVB_31_RIGHT | TGAGCCTTAATAGCTGCATCTGT |
| A1 | RSVA_33_LEFT | AAAAATCTGCTATCAAGAGTATGCCA | B1 | RSVB_33_LEFT | TGGACATCCAATGGTTGATGAAAGA |
| A1 | RSVA_33_RIGHT | TCCAACAAGGAAGGATAAGTGTTTAGT | B1 | RSVB_33_RIGHT | TGTGATGGCATGTAATTTCTAGGAAA |
| A1 | RSVA_35_LEFT | CTGATAGTGATAAATCAAGAAGAGTATTAGAGT | B1 | RSVB_35_LEFT | GCAACCAGGTATGTTTAGGCAAAT |
| A1 | RSVA_35_RIGHT | CGAAATGCTTGATTGAATTTGCTGA | B1 | RSVB_35_RIGHT | ACTTCATTAAGATTGACAACATGATCCT |
| A1 | RSVA_37_LEFT | AGGGTGGTGTCAAAAACTATGGA | B1 | RSVB_37_LEFT | CAGTCAATTGATATAAGTAAACCAGTTAGAC |
| A1 | RSVA_37_RIGHT | TCTTGTGTCAAACTACCTATAGATTCTAGA | B1 | RSVB_37_RIGHT | ACATAAAGCATGATTCCGGAGTTG |
| A1 | RSVA_39_LEFT | ATTTGCCCATGTTATTTGGTGGT | B1 | RSVB_39_LEFT | TGTTATATCGAAGCTTTTATAGGAGAACTC |
| A1 | RSVA_39_RIGHT | CTGCATAATATCATTAAGATCTATCTCTGTAGT | B1 | RSVB_39_RIGHT | TCATAAACAACTCTTAATCCATGAGGG |
| A1 | RSVA_41_LEFT | ACCATTAGATTGTAACAGAGATAAAAGAGA | B1 | RSVB_41_LEFT | AGCATAACTGAATTAAGCAAGTATGTAAGA |
| A1 | RSVA_41_RIGHT | AACCCAAGAGTTCCTATGCTAAGT | B1 | RSVB_41_RIGHT | TGTGGAAACAACTTTTTGGCCT |
| A1 | RSVA_43_LEFT | AGCTTTGGCCTTAGCTTAATGTCT | B1 | RSVB_43_LEFT | TCTTAGCTTGATGTCAGTTGTGGA |
| A1 | RSVA_43_RIGHT | TGATCAGTTATATATCCCTCTCCCCA | B1 | RSVB_43_RIGHT | TGAACATATGATCAGTTATATACCCCTCT |
| A1 | RSVA_45_LEFT | ACATAGAGTGAAGGGATGTCATAGC | B1 | RSVB_45_LEFT | AGTTTGCATAGAATAAAAGGTTGTCACA |
| A1 | RSVA_45_RIGHT | ACCTATACAATAGTCACTCAGTGTCTT | B1 | RSVB_45_RIGHT | CCACTTATACAAAATTTAGGTTTGTTTCTA |
| A1 | RSVA_47_LEFT | CTTCTCATCAAATATCCTTAGTGCACA | B1 | RSVB_47_LEFT | TGCATCACTTTATTGCATGCTTCC |
| A1 | RSVA_47_RIGHT | ACAAAAAGATTGATAGGTTCAGCAAACT | B1 | RSVB_47_RIGHT | ACAGGTAATTCAGCATCACAGACA |
| A1 | RSVA_49_LEFT | GGTCCTGCAAATGTGTTCCCA | B1 | RSVB_49_LEFT | ACTTCCTGTTTTTGATGTTGTGCA |
| A1 | RSVA_49_RIGHT | GGTTGTCAAGCTGTTTAACAATTCAC | B1 | RSVB_49_RIGHT | GCTTCTTGAGCTCATTGGTTGT |
| A2 | RSVA_2_LEFT | TGGCCTAATAGATGACAATTGTGAAA | B2 | RSVB_2_LEFT | ACAATGCCAATATTACAAAATGGAGGA |
| A2 | RSVA_2_RIGHT | TGACCAGAAATGTAAATGTGGCCT | B2 | RSVB_2_RIGHT | TGTGTGTGATGATTTCTTTGGTGAG |
| A2 | RSVA_4_LEFT | ACAACTTTATGCATAATCACACTCCA | B2 | RSVB_4_LEFT | GCATTAAGCCTACAAAACATACTCCT |
| A2 | RSVA_4_RIGHT | TGTTACATCCACTCCATTTGCCT | B2 | RSVB_4_RIGHT | TCTGGACATAGCATATAGCATACCTATT |
| A2 | RSVA_6_LEFT | AGCAGCAGGAGATAGATCAGGT | B2 | RSVB_6_LEFT | GCATGATTCTCCAGACTGTGGG |
| A2 | RSVA_6_RIGHT | TGCTTTTGGGTTGTTCAATATATGGT | B2 | RSVB_6_RIGHT | TTGCTCCATTTCTGCTTGGACA |
| A2 | RSVA_8_LEFT | TGACAGCAGAAGAACTAGAGGC | B2 | RSVB_8_LEFT | CAGAGCAACTCAAAGAAAATGGAGT |
| A2 | RSVA_8_RIGHT | TGAAAAAGGATTATCACTTGGCGT | B2 | RSVB_8_RIGHT | GCTTACTAGGGGTTTTCTTGGGT |
| A2 | RSVA_10_LEFT | GTGAAATACTAGGAATGCTTCACACA | B2 | RSVB_10_LEFT | TACTTCGGCTCGTGACGGAATA |
| A2 | RSVA_10_RIGHT | TTTTGGCTGGTTGGCTAATCAG | B2 | RSVB_10_RIGHT | TTGTCAGGGTGCTGATTGATCA |
| A2 | RSVA_12_LEFT | AGCCAATGTCAATATACTAGTGAAACAA | B2 | RSVB_12_LEFT | CAGATCTCTACGCCCAAAGGAC |
| A2 | RSVA_12_RIGHT | TGGCATTTTTGAATTCAGTGGTTGT | B2 | RSVB_12_RIGHT | AGCATTGGTGATAGCATTTTTGAATTC |
| A2 | RSVA_14_LEFT | ACATCAATGAGTAGATTCATACAAACTTTC | B2 | RSVB_14_LEFT | AGGTTCACATATATCCTCAACTGCA |
| A2 | RSVA_14_RIGHT | TGGAATACATTATATTCGCAGAGTTTGT | B2 | RSVB_14_RIGHT | GAGTTTTGTTGCAGAATATTTTATGTTCAC |
| A2 | RSVA_16_LEFT | GGGCAAATGCAAACATGTCCAA | B2 | RSVB_16_LEFT | GGGATCAAAAACAACATTGGGGC |
| A2 | RSVA_16_RIGHT | TTGTGGATTGTGGGGTTGACTC | B2 | RSVB_16_RIGHT | TTGGGTGATATTGTGGCTGAGT |
| A2 | RSVA_18_LEFT | ATCCAACCTGCTGGGCTATCT | B2 | RSVB_18_LEFT | ACCAACTACAAAACCCACAAACAA |
| A2 | RSVA_18_RIGHT | AGATTGTGATGGGTACTCGGATG | B2 | RSVB_18_RIGHT | ATCACGGTTCTCTGCTAAGATGT |
| A2 | RSVA_20_LEFT | TCCAGTCAAAACATCACTGAAGAATT | B2 | RSVB_20_LEFT | TTGGTATACCAGTGTCATAACAATAGAATT |
| A2 | RSVA_20_RIGHT | CAGGACCTTGGATACGGCAAT | B2 | RSVB_20_RIGHT | ACACTGACCCCATTTGATAGACTG |
| A2 | RSVA_22_LEFT | AGTTCCAACAAAAGAACAACAGACT | B2 | RSVB_22_LEFT | ACATGTTAACAAACAGTGAGTTACTATCA |
| A2 | RSVA_22_RIGHT | TGTTTCAGCTTGTGGGAAAAAGG | B2 | RSVB_22_RIGHT | AGTGTTACAAAGGCTGACTTCACT |
| A2 | RSVA_24_LEFT | ACAGCATCCAATAAAAATCGTGGG | B2 | RSVB_24_LEFT | ATGTCAACAAGCTGGAAGGCAA |
| A2 | RSVA_24_RIGHT | GTGCTTCTGGCCTTGCAGTATA | B2 | RSVB_24_RIGHT | TGGTCAGCAGATTGTTGTTGGA |
| A2 | RSVA_26_LEFT | AACCATCTCACTTACACTTTTTAAGTAGA | B2 | RSVB_26_LEFT | TGCTTGAATGGTAGAAGATGCCA |
| A2 | RSVA_26_RIGHT | CAGTAAGGAGTTTGCTCATGGC | B2 | RSVB_26_RIGHT | AGATGGATGGTTTGCTTGCTGT |
| A2 | RSVA_28_LEFT | AGCATAACCATCAATAACCCAAAAGA | B2 | RSVB_28_LEFT | AACCACAACCATTTAGATAACCACC |
| A2 | RSVA_28_RIGHT | AGCAGAATTTCCGCTAATAATGGGA | B2 | RSVB_28_RIGHT | TGCTCTATTAGTGGGCTTTGTCT |
| A2 | RSVA_30_LEFT | GTCCTTAATATCCAAGTATCATAAAGGTGA | B2 | RSVB_30_LEFT | GGAAAAGGACAGAGTTAAGCCCA |
| A2 | RSVA_30_RIGHT | AACCATGATGGAGGATGTTGCA | B2 | RSVB_30_RIGHT | GTCAAAAATTGATTGTATGTGGTAGTTGT |
| A2 | RSVA_32_LEFT | TCTTGACATGGAAAGATATTAGCCTTAGT | B2 | RSVB_32_LEFT | GAAGGCTTTTACATAATAAAAGAAGTAGAAGG |
| A2 | RSVA_32_RIGHT | ACTTACTCAATAGAATTATCCATCTGCCA | B2 | RSVB_32_RIGHT | GCACCTCTCAACGTACTTAGGC |
| A2 | RSVA_34_LEFT | ACAGATGGCCTACTTTAAGAAATGC | B2 | RSVB_34_LEFT | TCTATCGTGAGTTTCATCTGCCT |
| A2 | RSVA_34_RIGHT | ACATGATTCGGGTTGTTAAGATAACTT | B2 | RSVB_34_RIGHT | GTCAAACTCTCAGGGAAGAATTGTAA |
| A2 | RSVA_36_LEFT | AGCAGGAATAAGTAACAAATCAAATCGT | B2 | RSVB_36_LEFT | TGGTTGCATTTAACAATACCTCTTGT |
| A2 | RSVA_36_RIGHT | TGTCACCATTAATTAAAGCAGTAATTGAGA | B2 | RSVB_36_RIGHT | TCGAGATATATAGGTCTCAGTTCCCT |
| A2 | RSVA_38_LEFT | ACCCAGCTAGTATAAAGAAAGTCCTAA | B2 | RSVB_38_LEFT | ACACAGGAGTTAGAATACAGAGGAGA |
| A2 | RSVA_38_RIGHT | TGAACACAGAGTGAACTATAGCCT | B2 | RSVB_38_RIGHT | ACACAAGTCAAGAATTTGTTCAGTCT |
| A2 | RSVA_40_LEFT | TAGACTGGCAGTTACCGAGGTT | B2 | RSVB_40_LEFT | GTGCGCAACACTATACTACCACT |
| A2 | RSVA_40_RIGHT | GGATAAAGACCAAGATCTTTCTCTAACG | B2 | RSVB_40_RIGHT | TGTCCATTGTGAACATAATACTTGGTG |
| A2 | RSVA_42_LEFT | TCTATTAGCAAAATTGGATTGGGTGT | B2 | RSVB_42_LEFT | TGCATCCATAGACAACAAAGATGAAT |
| A2 | RSVA_42_RIGHT | TGGGAGGTTTCATCAAATGTATCTCA | B2 | RSVB_42_RIGHT | TCAACTTGATGATATCAACATCTCCTG |
| A2 | RSVA_44_LEFT | ACTAATTTAGCTGGACATTGGATTCTT | B2 | RSVB_44_LEFT | GCTGGACATTGGATTCTGATTATTCA |
| A2 | RSVA_44_RIGHT | TAACAACCCAAGGGCAAACTGT | B2 | RSVB_44_RIGHT | GTTAACAACCCAAGGGCATACG |
| A2 | RSVA_46_LEFT | AGGATTGCTAATTCCGAATTAGAAAGT | B2 | RSVB_46_LEFT | ATATCACCCAACCCCAGAAGCT |
| A2 | RSVA_46_RIGHT | CAACCTGTAGAACTAAATACAAAATTGAATCTG | B2 | RSVB_46_RIGHT | ATATTCTATACTGATCTTGCATCCTGTG |
| A2 | RSVA_48_LEFT | GGTGAAAATTTGACCATTCCTGCTA | B2 | RSVB_48_LEFT | TCCTGCTACAGATGCAACTAACA |
| A2 | RSVA_48_RIGHT | GGCATGATGAAATTTTTGGTTCTTGA | B2 | RSVB_48_RIGHT | AGATTCCTTGTCAATCTTTTTAGGCATA |
| A2 | RSVA_50_LEFT | AGAGTGTTGTTAGTGGAGATATACTATCA | B2 | RSVB_50_LEFT | GCAACAAGCTTATAAACCACAAGC |
| A2 | RSVA_50_RIGHT | CCTAGATCAAAATGATAATTTTAGGATTGGTTC | B2 | RSVB_50_RIGHT | TGTCTCGTTGTGTTGTAAATGCAC |

Table S2. PCR primer sequences

| Primer/Probe Name | Primer/Probe Sequence |
| --- | --- |
| RSV Typing Forward Primer | AAT ACA GCC AAA TCT AAC CAA CTT TAC A |
| RSV Typing Reverse Primer | GCC AAG GAA GCA TGC AAT AAA |
| RSV A Typing Probe | TGC TAY TGT GCA CTA AAG |
| RSV B Typing Probe | CAC TAT TCC TTA CTA AAG ATG TC |
| EXO3 Forward Primer | AAT TGG AAG TGG CGG AAG AA |
| EXO3 Reverse Primer | GGA ACC TAA GAC AAG TGT GTT TAT GG |
| EXO3 Probe | Cy5-AGC TAT TGC AAA CGC CAT CGC ACA A-BHQ2 |
| RSV Quantification Forward Primer 1 | GGA AAC ATA CGT GAA YAA ACT TCA |
| RSV Quantification Forward Primer 2 | GGA AAC ATA CGT GAA CAA GCT TCA |
| RSV Quantification Reverse Primer 1 | CAT CGT CTT TTT CTA RGA CAT TGT ATT GA |
| RSV Quantification Reverse Primer 2 | TCA TCA TCT TTT TCT AGA ACA TTG TAC TGA |
| RSV Quantification Probe | 6FAM-TGT GTA TGT GGA GCC YT-NFQMGB |
| EXO2 Forward Primer | GGC GGA AGA ACA GCT ATT GC |
| EXO2 Reverse Primer | GGA ACC TAA GAC AAG TGT GTT TAT GG |
| EXO2 Probe | VIC-AAC GCC ATC GCA CAA T-NGQMGB |

Table S3. Sanger sequencing primers

| Sanger sequencing Primer Name | Sequence |
| --- | --- |
| RSVA-F Fwd2 (F2) | GCTCACCTCCAACACCAAAG |
| RSVA-F Rev1 (R1) | CATGGGGTGGCCATTCAAAA |
| RSVA-F SeqA R1 | GCTGGTTAAGACACTGACTCC |
| RSVA-F SeqA R2 | AGTCACACCTGCATTAACACT |
| RSVA-F SeqB F1 | AGGTGTTGGATCTGCAATCG |
| RSVA-F SeqB R1 | ACGGAGCTGCTTACATCTGT |
| RSVA-F SeqB R2 | AGCATGACACAATGGCTCCT |
| RSVA-F SeqC F1 | TGGAAACTACACACATCCCCT |
| RSVA-F SeqC F2 | TCCCACAAGCTGAAACATGT |
| RSVA-F SeqC R1 | GGTCAAATAGCGAACCATTGTAA |
| RSVA-F SeqC R2 | CGTGACATATTTGCCCCAGT |

Table S4. Specificity of UW-ARTIC RSV amplicon panel

|  | **RSVA** | | | | | | | **RSVB** | | | | | | |
| --- | --- | --- | --- | --- | --- | --- | --- | --- | --- | --- | --- | --- | --- | --- |
| **Sample** | **raw reads** | **mapped reads** | **pct reads mapped** | **mean genome coverage** | **mean fusion coverage** | **pct fusion 100x** | **%Ns** | **raw reads** | **mapped reads** | **pct reads mapped** | **mean genome coverage** | **mean fusion coverage** | **pct fusion 100x** | **%Ns** |
| RSV_Spec_01 | 5641240 | 66 | 0.00 | 0.16 | 0.15 | 0 | 99.23 | 2347662 | 490 | 0.02 | 1.04 | 0.03 | 0 | 99.78 |
| RSV_Spec_02 | 3107496 | 53 | 0.00 | 0.12 | 0.48 | 0 | 99.77 | 3068806 | 571 | 0.02 | 1.22 | 0.18 | 0 | 99.17 |
| RSV_Spec_03 | 3280028 | 37 | 0.00 | 0.08 | 0.00 | 0 | 99.76 | 6432670 | 482 | 0.01 | 1.02 | 0.03 | 0 | 99.17 |
| RSV_Spec_04 | 4477954 | 30 | 0.00 | 0.06 | 0.00 | 0 | 99.80 | 2631540 | 207 | 0.01 | 0.44 | 0.03 | 0 | 99.58 |
| RSV_Spec_05 | 9414838 | 39 | 0.00 | 0.12 | 0.07 | 0 | 99.80 | 9111276 | 261 | 0.00 | 0.59 | 0.16 | 0 | 99.58 |
| RSV_Spec_06 | 6850554 | 40 | 0.00 | 0.10 | 0.28 | 0 | 99.62 | 9415890 | 317 | 0.00 | 0.69 | 0.18 | 0 | 99.15 |
| RSV_Spec_07 | 2784894 | 42 | 0.00 | 0.09 | 0.04 | 0 | 99.56 | 4544324 | 447 | 0.01 | 0.95 | 0.00 | 0 | 99.58 |
| RSV_Spec_08 | 5093360 | 25 | 0.00 | 0.06 | 0.11 | 0 | 99.42 | 8257212 | 331 | 0.00 | 0.73 | 0.00 | 0 | 99.56 |
| RSV_Spec_09 | 1251920 | 38 | 0.00 | 0.09 | 0.34 | 0 | 99.59 | 4594392 | 412 | 0.01 | 0.91 | 0.07 | 0 | 99.39 |
| RSV_Spec_10 | 860748 | 46 | 0.01 | 0.10 | 0.29 | 0 | 99.57 | 2052204 | 475 | 0.02 | 1.01 | 0.10 | 0 | 99.58 |
| RSV_Spec_11 | 977364 | 53 | 0.01 | 0.12 | 0.30 | 0 | 99.35 | 1073882 | 532 | 0.05 | 1.13 | 0.00 | 0 | 99.38 |
| RSV_Spec_12 | 458220 | 84 | 0.02 | 0.18 | 0.59 | 0 | 98.93 | N/A | N/A | N/A | N/A | N/A | N/A | N/A |
| RSV_Spec_13 | 4677660 | 28 | 0.00 | 0.06 | 0.08 | 0 | 99.80 | N/A | N/A | N/A | N/A | N/A | N/A | N/A |
| RSV_Spec_14 | 481430 | 53 | 0.01 | 0.12 | 0.31 | 0 | 99.17 | N/A | N/A | N/A | N/A | N/A | N/A | N/A |
| RSV_Spec_15 | 408910 | 67 | 0.02 | 0.15 | 0.41 | 0 | 99.55 | 417186 | 1100 | 0.26 | 2.34 | 0.07 | 0 | 99.78 |
| RSV_Spec_16 | 328208 | 36 | 0.01 | 0.08 | 0.26 | 0 | 99.36 | 231118 | 771 | 0.33 | 1.64 | 0.04 | 0 | 99.57 |
| RSV_Spec_17 | 3644116 | 76 | 0.00 | 0.18 | 0.22 | 0 | 98.66 | 4826446 | 523 | 0.01 | 1.13 | 0.00 | 0 | 99.78 |
| RSV_Spec_18 | 2634414 | 13 | 0.00 | 0.03 | 0.08 | 0 | 99.80 | 10564276 | 507 | 0.00 | 1.10 | 0.04 | 0 | 99.58 |
| RSV_Spec_19 | 5312964 | 24 | 0.00 | 0.05 | 0.00 | 0 | 99.57 | 8984836 | 342 | 0.00 | 0.74 | 0.00 | 0 | 99.58 |
| RSV_Spec_20 | 343152 | 49 | 0.01 | 0.11 | 0.41 | 0 | 99.13 | 774376 | 638 | 0.08 | 1.36 | 0.00 | 0 | 99.57 |
| RSV_Spec_21 | 2350260 | 33 | 0.00 | 0.08 | 0.11 | 0 | 99.60 | 4112170 | 380 | 0.01 | 0.81 | 0.00 | 0 | 99.58 |
| RSV_Spec_22 | 674784 | 82 | 0.01 | 0.19 | 0.40 | 0 | 99.33 | 1058238 | 779 | 0.07 | 1.66 | 0.03 | 0 | 99.57 |
| RSV_Spec_23 | 636302 | 172 | 0.03 | 0.38 | 0.49 | 0 | 99.32 | 245316 | 692 | 0.28 | 1.47 | 0.00 | 0 | 99.57 |
| RSV_Spec_24 | 840952 | 67 | 0.01 | 0.14 | 0.45 | 0 | 99.17 | N/A | N/A | N/A | N/A | N/A | N/A | N/A |
| RSV_Spec_25 | 1953988 | 69 | 0.00 | 0.15 | 0.28 | 0 | 99.08 | 3405648 | 608 | 0.02 | 1.29 | 0.00 | 0 | 99.58 |
| RSV_Spec_26 | 4075602 | 39 | 0.00 | 0.13 | 0.00 | 0 | 99.59 | 5189786 | 388 | 0.01 | 0.84 | 0.09 | 0 | 99.38 |
| RSV_Spec_27 | 4663692 | 25 | 0.00 | 0.06 | 0.19 | 0 | 99.79 | 7064532 | 471 | 0.01 | 1.00 | 0.03 | 0 | 99.38 |
| RSV_Spec_28 | 3579680 | 46 | 0.00 | 0.11 | 0.22 | 0 | 99.35 | 6379500 | 486 | 0.01 | 1.06 | 0.00 | 0 | 98.96 |
| RSV_Spec_29 | 2039136 | 39 | 0.00 | 0.09 | 0.14 | 0 | 99.59 | 3631436 | 563 | 0.02 | 1.20 | 0.00 | 0 | 99.37 |
| RSV_Spec_30 | 2004150 | 43 | 0.00 | 0.09 | 0.26 | 0 | 99.55 | 4273282 | 437 | 0.01 | 0.96 | 0.18 | 0 | 99.59 |
| RSV_Spec_31 | 5232566 | 35 | 0.00 | 0.08 | 0.15 | 0 | 99.39 | 2797622 | 98 | 0.00 | 0.21 | 0.00 | 0 | 99.38 |
| RSV_Spec_32 | 4859056 | 18 | 0.00 | 0.04 | 0.04 | 0 | 99.57 | 8437596 | 400 | 0.00 | 0.85 | 0.07 | 0 | 98.98 |
| RSV_Spec_33 | 3484618 | 46 | 0.00 | 0.11 | 0.22 | 0 | 99.35 | 5857396 | 450 | 0.01 | 0.97 | 0.11 | 0 | 99.58 |
| RSV_Spec_34 | 1135454 | 30 | 0.00 | 0.06 | 0.19 | 0 | 99.38 | 3289312 | 464 | 0.01 | 0.98 | 0.07 | 0 | 99.59 |
| RSV_Spec_35 | 2374570 | 86 | 0.00 | 0.19 | 0.44 | 0 | 98.96 | 2473274 | 734 | 0.03 | 1.59 | 0.27 | 0 | 99.58 |
| RSV_Spec_36 | 7342232 | 35 | 0.00 | 0.12 | 0.28 | 0 | 99.35 | 10861498 | 193 | 0.00 | 0.43 | 0.00 | 0 | 99.77 |
| RSV_Spec_37 | 2497932 | 11 | 0.00 | 0.03 | 0.00 | 0 | 100.00 | 4415866 | 338 | 0.01 | 0.75 | 0.42 | 0 | 99.59 |
| RSV_Spec_38 | 7178572 | 45 | 0.00 | 0.11 | 0.11 | 0 | 99.16 | 7453236 | 146 | 0.00 | 0.34 | 0.09 | 0 | 99.58 |
| RSV_Spec_39 | 620550 | 100 | 0.02 | 0.22 | 0.50 | 0 | 98.76 | 448558 | 689 | 0.15 | 1.46 | 0.00 | 0 | 99.59 |
| RSV_Spec_40 | 6236744 | 42 | 0.00 | 0.12 | 0.11 | 0 | 99.60 | 2591252 | 441 | 0.02 | 1.00 | 0.17 | 0 | 99.38 |

Table S5. Clinical samples used to test breadth of genome recovery

| **Specimen** | **Copies/mL** | **Strain name** | **Collection date** | **GenBank** | **BioSample** | **SRA** | **Lineage** |
| --- | --- | --- | --- | --- | --- | --- | --- |
| RSV-A 01 | 26700000 | RSV/A/USA/WA-UW-05541/2022 | 2022-10 | PP760388 | SAMN41195160 | SRR28891439 | A.D.5.2 |
| RSV-A 02 | 2920000 | RSV/A/USA/WA-UW-05560/2022 | 2022-10 | PP760389 | SAMN41195161 | SRR28891438 | A.D.3 |
| RSV-A 03 | 15800000 | RSV/A/USA/WA-UW-05648/2022 | 2022-10 | PP760392 | SAMN41195162 | SRR28891427 | A.D.5.2 |
| RSV-A 04 | 2570000 | RSV/A/USA/WA-UW-05654/2022 | 2022-10 | PP760393 | SAMN41195163 | SRR28891419 | A.D.3.1 |
| RSV-A 05 | 2680000 | RSV/A/USA/WA-UW-05680/2022 | 2022-11 | PP760396 | SAMN41195164 | SRR28891418 | A.D.5.2 |
| RSV-A 06 | 988000 | RSV/A/USA/WA-UW-05694/2022 | 2022-11 | PP760397 | SAMN41195165 | SRR28891417 | A.D.3 |
| RSV-A 07 | 30000000 | RSV/A/USA/WA-UW-05700/2022 | 2022-11 | PP760398 | SAMN41195166 | SRR28891416 | A.D.5.2 |
| RSV-A 08 | 1030000 | RSV/A/USA/WA-UW-05704/2022 | 2022-11 | PP760399 | SAMN41195167 | SRR28891415 | A.D.5.2 |
| RSV-A 09 | 8090000 | RSV/A/USA/WA-UW-05708/2022 | 2022-11 | PP760401 | SAMN41195168 | SRR28891414 | A.D.5.2 |
| RSV-A 10 | 1180000 | RSV/A/USA/WA-UW-R3AE2/2022 | 2022-10 | PP760419 | SAMN32118081 | SRR28895551 | A.D.1.4 |
| RSV-A 11 | 1290000 | RSV/A/USA/WA-UW-3TBF2/2022 | 2022-10 | PP760385 | SAMN32118086 | SRR28895550 | A.D.5.2 |
| RSV-A 12 | 7860000 | RSV/A/USA/WA-UW-34AM6/2022 | 2022-11 | PP760387 | SAMN32118092 | SRR28895546 | A.D.5.2 |
| RSV-A 13 | 13700000 | RSV/A/USA/WA-UW-TQVY1/2022 | 2022-11 | PP760422 | SAMN32118087 | SRR28895545 | A.D.3 |
| RSV-A 14 | 2010000 | RSV/A/USA/WA-UW-DT79D/2022 | 2022-10 | PP760416 | SAMN32118090 | SRR28895544 | A.D.5.2 |
| RSV-A 15 | 40100000 | RSV/A/USA/WA-UW-19667/2023 | 2023-01 | PP760406 | SAMN41195169 | SRR28891413 | A.D.3 |
| RSV-A 16 | 165000 | RSV/A/USA/WA-UW-71432/2022 | 2022-12 | PP760412 | SAMN41195170 | SRR28891437 | A.D.5.2 |
| RSV-A 17 | 67500000 | RSV/A/USA/WA-UW-15500/2022 | 2022-12 | PP760404 | SAMN41195171 | SRR28891436 | A.D.5.2 |
| RSV-A 18 | 5340 | N/A | 2023-02 | N/A | N/A | N/A | N/A |
| RSV-A 19 | 7740 | RSV/A/USA/WA-UW-32155/2022 | 2023-01 | PP760408 | SAMN41195173 | SRR28891434 | A.D.5.2 |
| RSV-A 20 | 21800000 | RSV/A/USA/WA-UW-94220/2022 | 2022-12 | PP760414 | SAMN41195174 | SRR28891433 | A.D.5 |
| RSV-B 01 | 238000 | RSV/B/USA/WA-UW-05596/2022 | 2022-10 | PP760390 | SAMN41195175 | SRR28891432 | B.D.E.1 |
| RSV-B 02 | 137000 | RSV/B/USA/WA-UW-05677/2022 | 2022-11 | PP760395 | SAMN41195178 | SRR28891429 | B.D.E.1 |
| RSV-B 03 | 196000 | RSV/B/USA/WA-UW-05662/2022 | 2022-10 | PP760394 | SAMN41195177 | SRR28891430 | B.D.E.1 |
| RSV-B 04 | 226000 | RSV/B/USA/WA-UW-05631/2022 | 2022-10 | PP760391 | SAMN41195176 | SRR28891431 | B.D.E.1 |
| RSV-B 05 | 38700 | RSV/B/USA/WA-UW-05706/2022 | 2022-11 | PP760400 | SAMN41195179 | SRR28891428 | B.D.E.1 |
| RSV-B 06 | 7000000 | RSV/B/USA/WA-UW-KH4F2/2022 | 2022-10 | PP760417 | SAMN32118096 | SRR28895543 | B.D.E.1 |
| RSV-B 07 | 5590000 | RSV/B/USA/WA-UW-RFHA8/2022 | 2022-11 | PP760420 | SAMN32118103 | SRR28895542 | B.D.E.1 |
| RSV-B 08 | 2160000 | RSV/B/USA/WA-UW-YFP3F/2022 | 2022-10 | PP760423 | SAMN32118116 | SRR28895541 | B.D.E.1 |
| RSV-B 09 | 4100000 | RSV/B/USA/WA-UW-M4SWF/2022 | 2022-10 | PP760418 | SAMN32118117 | SRR28895540 | B.D.E.1 |
| RSV-B 10 | 15300000 | RSV/B/USA/WA-UW-RWWYB/2022 | 2022-10 | PP760421 | SAMN32118097 | SRR28895539 | B.D.E.1 |
| RSV-B 11 | 639000 | RSV/B/USA/WA-UW-YVED3/2022 | 2022-11 | PP760424 | SAMN32118113 | SRR28895549 | B.D.E.1 |
| RSV-B 12 | 362000 | RSV/B/USA/WA-UW-A7421/2022 | 2022-10 | PP760415 | SAMN32118094 | SRR28895548 | B.D.E.1 |
| RSV-B 14 | 5100 | RSV/B/USA/WA-UW-09863/2023 | 2023-01 | PP760403 | SAMN41195185 | SRR28891421 | B.D.E.1 |
| RSV-B 13 | 540000 | RSV/B/USA/WA-UW-8BLN9/2022 | 2022-10 | PP760386 | SAMN32118102 | SRR28895547 | B.D.E.1 |
| RSV-B 15 | 3980 | RSV/B/USA/WA-UW-06989/2022 | 2022-12 | PP760402 | SAMN41195181 | SRR28891425 | B.D.E.1 |
| RSV-B 16 | 10900 | RSV/B/USA/WA-UW-18337/2022 | 2022-12 | PP760405 | SAMN41195182 | SRR28891424 | B.D.E.1 |
| RSV-B 17 | 476000 | RSV/B/USA/WA-UW-68476/2022 | 2022-12 | PP760410 | SAMN41195183 | SRR28891423 | B.D.E.1 |
| RSV-B 18 | 674000 | RSV/B/USA/WA-UW-69563/2023 | 2023-02 | PP760411 | SAMN41195186 | SRR28891420 | B.D.E.1 |
| RSV-B 19 | 197000 | RSV/B/USA/WA-UW-25072/2022 | 2022-12 | PP760407 | SAMN41195180 | SRR28891426 | B.D.E.1 |
| RSV-B 20 | 4360000 | RSV/B/USA/WA-UW-65453/2023 | 2023-01 | PP760409 | SAMN41195184 | SRR28891422 | B.D.E.1 |

Table S6. Reduced amplification resulting in full (200bp or greater stretches of Ns) or partial (100-200 Ns) amplicon dropout

| Insert | Full | Partial |
| --- | --- | --- |
| RSVA_3 | 3 | 1 |
| RSVA_4 | 1 | 0 |
| RSVA_13 | 3 | 1 |
| RSVA_33 | 1 | 0 |
| RSVA_45 | 0 | 1 |
| RSVA_47 | 5 | 2 |
| RSVB_5 | 0 | 2 |
| RSVB_11 | 1 | 1 |
| RSVB_13 | 0 | 1 |
| RSVB_17 | 1 | 0 |
| RSVB_19 | 1 | 0 |
| RSVB_25 | 1 | 0 |
| RSVB_29 | 3 | 1 |
| RSVB_31 | 1 | 1 |
| RSVB_37 | 0 | 1 |
| RSVB_39 | 0 | 1 |
| RSVB_41 | 1 | 2 |
| RSVB_43 | 1 | 0 |
| RSVB_45 | 3 | 0 |
| RSVB_47 | 1 | 1 |
| RSVB_49 | 0 | 3 |

Table S7. Consensus-level differences between samples prepared twice by the same technician and sequenced on the same run (repeatability/intra-run precision)

| **Specimen** | **Pairwise differences** (before masking Ns) | **Positions with Ns** | **Pairwise differences**  (after masking Ns) |
| --- | --- | --- | --- |
| RSV-A 01 | 189 | 1248 | 0 |
| RSV-A 02 | 413 | 1706 | 2 |
| RSV-A 03 | 150 | 1271 | 0 |
| RSV-A 04 | 0 | 1687 | 0 |
| RSV-A 05 | 402 | 1404 | 0 |
| RSV-A 06 | 53 | 920 | 0 |
| RSV-A 07 | 259 | 931 | 0 |
| RSV-A 08 | 425 | 1639 | 0 |
| RSV-A 09 | 242 | 952 | 0 |
| RSV-A 10 | 11 | 424 | 0 |
| RSV-A 11 | 68 | 1755 | 0 |
| RSV-A 12 | 160 | 1646 | 0 |
| RSV-A 13 | 264 | 1102 | 0 |
| RSV-A 14 | 48 | 1635 | 1 |
| RSV-A 15 | 215 | 878 | 0 |
| RSV-A 16 | 172 | 1648 | 1 |
| RSV-A 17 | 179 | 829 | 0 |
| RSV-A 20 | 23 | 410 | 0 |
| RSV-B 03 | 367 | 544 | 1 |
| RSV-B 06 | 803 | 937 | 1 |
| RSV-B 07 | 271 | 406 | 0 |
| RSV-B 08 | 246 | 380 | 1 |
| RSV-B 11 | 0 | 135 | 0 |
| RSV-B 17 | 0 | 135 | 0 |
| RSV-B 18 | 0 | 135 | 0 |
| RSV-B 20 | 155 | 307 | 0 |

Table S7. Consensus-level differences between samples prepared twice by different technicians and sequenced on different runs (reproducibility/inter-run precision)

| **Specimen** | **Pairwise differences** (before masking Ns) | **Positions with Ns** | **Pairwise differences**  (after masking Ns) |
| --- | --- | --- | --- |
| RSV-A 01 | 731 | 1248 | 0 |
| RSV-A 02 | 1455 | 1728 | 2 |
| RSV-A 03 | 476 | 1271 | 0 |
| RSV-A 04 | 1047 | 1687 | 0 |
| RSV-A 05 | 823 | 1404 | 0 |
| RSV-A 06 | 537 | 920 | 0 |
| RSV-A 07 | 474 | 950 | 0 |
| RSV-A 08 | 1315 | 1639 | 0 |
| RSV-A 09 | 735 | 952 | 0 |
| RSV-A 10 | 261 | 424 | 0 |
| RSV-A 12 | 869 | 1646 | 0 |
| RSV-A 13 | 699 | 1102 | 1 |
| RSV-A 14 | 581 | 1738 | 0 |
| RSV-A 15 | 741 | 878 | 0 |
| RSV-A 16 | 772 | 1648 | 2 |
| RSV-B 06 | 803 | 937 | 1 |
| RSV-B 07 | 271 | 406 | 0 |
| RSV-B 08 | 246 | 380 | 1 |
| RSV-B 11 | 0 | 135 | 0 |
| RSV-B 17 | 0 | 135 | 0 |
| RSV-B 18 | 0 | 135 | 0 |
| RSV-B 20 | 155 | 307 | 0 |
